# Chaperone Requirements for De Novo Folding of *Saccharomyces cerevisiae* Septins

**DOI:** 10.1101/2022.03.31.486562

**Authors:** Daniel Hassell, Ashley Denney, Emily Singer, Aleyna Benson, Andrew Roth, Julia Ceglowski, Marc Steingesser, Michael McMurray

## Abstract

Polymers of septin protein complexes play cytoskeletal roles in eukaryotic cells. The specific subunit composition within complexes controls functions and higher-order structural properties. All septins have globular GTPase domains. The other eukaryotic cytoskeletal NTPases strictly require assistance from molecular chaperones of the cytosol, particularly the cage-like chaperonins, to fold into oligomerization-competent conformations. We previously identified cytosolic chaperones that bind septins and influence the oligomerization ability of septins carrying mutations linked to human disease, but it was unknown to what extent wild-type septins require chaperone assistance for their native folding. Here we use a combination of *in vivo* and *in vitro* approaches to demonstrate chaperone requirements for *de novo* folding and complex assembly by budding yeast septins. Individually purified septins adopted non-native conformations and formed non-native homodimers. In chaperonin- or Hsp70-deficient cells, septins folded slower and were unable to assemble post-translationally into native complexes. One septin, Cdc12, was so dependent on co-translational chaperonin assistance that translation failed without it. Our findings point to distinct translation elongation rates for different septins as a possible mechanism to direct a stepwise, co-translational assembly pathway in which general cytosolic chaperones act as key intermediaries.

## INTRODUCTION

Septins evolved from ancient GTPases into a diverse family of eukaryotic proteins capable of assembling into hetero-oligomers (Shuman and Momany, 2021). Septin hetero-oligomers further polymerize into cytoskeletal filaments involved in a variety of cellular functions (Woods and Gladfelter, 2021). In many cell types no septin monomer can be found (Abbey et al., 2016; Bridges et al., 2014; Frazier et al., 1998). Similarly, the building blocks of microtubules are stable heterodimers of α and β tubulins, and individual tubulin proteins exist fleetingly, in complexes with cytosolic chaperones and dedicated folding co-factors (Lewis et al., 1997). The human septin mutation SEPT12(Asp197Asn) causes disease by interfering with filament assembly (Kuo et al., 2012) but was first identified 40 years prior in the homologous septin in the budding yeast *Saccharomyces cerevisiae*, as Cdc10(Asp182Asn) (Hartwell, 1971; McMurray et al., 2011a). We found that this mutation slows folding of the yeast septin and prolongs interactions with cytosolic chaperones, leading to a kinetic delay in the ability of the mutant septin to stably incorporate into functional septin hetero-oligomers (Denney et al., 2021; Johnson et al., 2015). Our other studies provided evidence that conformational changes at septin-septin interfaces – some linked to GTP binding and/or hydrolysis – are important for an efficient, step-wise pathway of *de novo* septin hetero-oligomer assembly (Johnson et al., 2015; Weems and McMurray, 2017; Weems et al., 2014). These findings support a model in which a newly translated septin can explore a variety of conformations, at least some of which bind cytosolic chaperones, before achieving its native conformation in the context of a septin hetero-oligomer. It was unclear to what extent chaperone function promotes wild-type septin folding and hetero-oligomerization.

Chaperonins are large, oligomeric chaperones that form a client-binding chamber and couple the hydrolysis of ATP to conformational changes that promote client folding and eventual release (Gestaut et al., 2019; Hayer-Hartl et al., 2016). Chaperonins diverged during evolution into two groups. A proteomics study querying the proteins bound by the Group II chaperonin of the *S. cerevisiae* cytosol –– called chaperonin containing tailless complex polypeptide 1 (CCT) or tailless complex polypeptide 1 ring complex (TRiC) –– and released upon ATP hydrolysis identified Cdc3, Cdc10, Cdc11, Cdc12 and Shs1, the five septin proteins that are expressed in mitotically dividing cells, (Dekker et al., 2008). An independent study also found co-purification with CCT of an overexpressed, GST-tagged yeast septin, Cdc10 (Yam et al., 2008). A more recent study specifically investigating co-translational CCT interactions identified ribosomes translating yeast septins (Stein et al., 2019). Finally, our split-protein complementation assay studies demonstrated in living yeast cells that multiple CCT subunits interact with multiple septins (Denney et al., 2021). Thus all available evidence supports direct interactions between nascent yeast septins and CCT. However, a role for CCT–septin interactions in promoting efficient *de novo* septin folding was previously dismissed (Dekker et al., 2008), for two reasons. First, septins successfully fold and hetero-oligomerize when heterologously expressed in *E. coli* (Bertin et al., 2008; Farkasovsky et al., 2005; Versele and Thorner, 2004; Versele et al., 2004), which lacks CCT and instead has a Group I chaperonin, GroEL. Second, yeast Cdc10 synthesized by *E. coli* ribosomes in a cell-free translation reaction migrates in native PAGE in a way that is unaffected by the addition of purified CCT (Dekker et al., 2008), suggesting that CCT does not influence Cdc10 conformation. Here we explore in greater detail chaperonin requirements for efficient septin synthesis and folding.

By “shielding” exposed regions normally buried in the core of a folded protein or in a protein-protein interface, chaperone binding helps client proteins avoid making inappropriate inter- and intra-molecular interactions and becoming “trapped” in low-energy, non-native conformations (Balchin et al., 2016). Septin-septin interactions in the context of native septin oligomers presumably also help the individual proteins maintain native conformations and avoid inappropriate interactions. Indeed, a purified single human septin is prone to amyloid aggregation under conditions where a septin heterodimer is not, suggesting that native septin oligomerization prevents amyloid formation (Kumagai et al., 2019). Similarly, early studies of septin subunit organization within hetero-oligomers were misled by non-native interactions found when individual septins were expressed individually in *E. coli* and purified away from all other proteins (including chaperones) prior to mixing with other individually expressed and purified septins (Versele et al., 2004). For example, purified Cdc12 interacted robustly with Cdc10 and with itself (Versele et al., 2004), but neither of these interactions occurs within native hetero-oligomers, which are linear octamers with subunit order Cdc11–Cdc12–Cdc3–Cdc10–Cdc10–Cdc3–Cdc12–Cdc11 (Bertin et al., 2008) or Shs1–Cdc12– Cdc3–Cdc10–Cdc10–Cdc3–Cdc12–Cdc11 (Garcia et al., 2011). Here we used genetic, biochemical and biophysical methods to obtain new insights into the conformations of “isolated” septins, their ability to assemble into native hetero-oligomers, and roles for cytosolic chaperones in maintaining that ability.

## RESULTS

### Non-native conformations and oligomerization by individual purified septins

To explore in greater detail the ways that yeast septins fold and oligomerize in the absence of chaperones and other septins, we used a published plasmid (Versele and Thorner, 2004) to purify hexahistidine-tagged (“6xHis”) Cdc3 following *E. coli* expression. In the original study, the oligomerization status of 6xHis-Cdc3 was not determined. It was capable of binding robustly to purified GST-Cdc12 or GST-Cdc10, the native interaction partners (Versele et al., 2004), suggesting that at least some molecules had interfaces available for native interactions. We used a centrifugal filtration column with a nominal 100-kDa molecular weight cutoff, and found that, like homotetrameric alcohol dehydrogenase (Adh1, native molecular weight ∼150 kDa, monomer molecular weight 37 kDa), 6xHis-Cdc3 (monomeric molecular weight ∼60 kDa) remained in the retentate fraction (Figure 1A), suggestive of homo-oligomerization. Consistent with all prior reports, Cdc3 migrated anomalously during SDS-PAGE, at an apparent molecular weight of ∼70 kDa.

**Figure 1.**
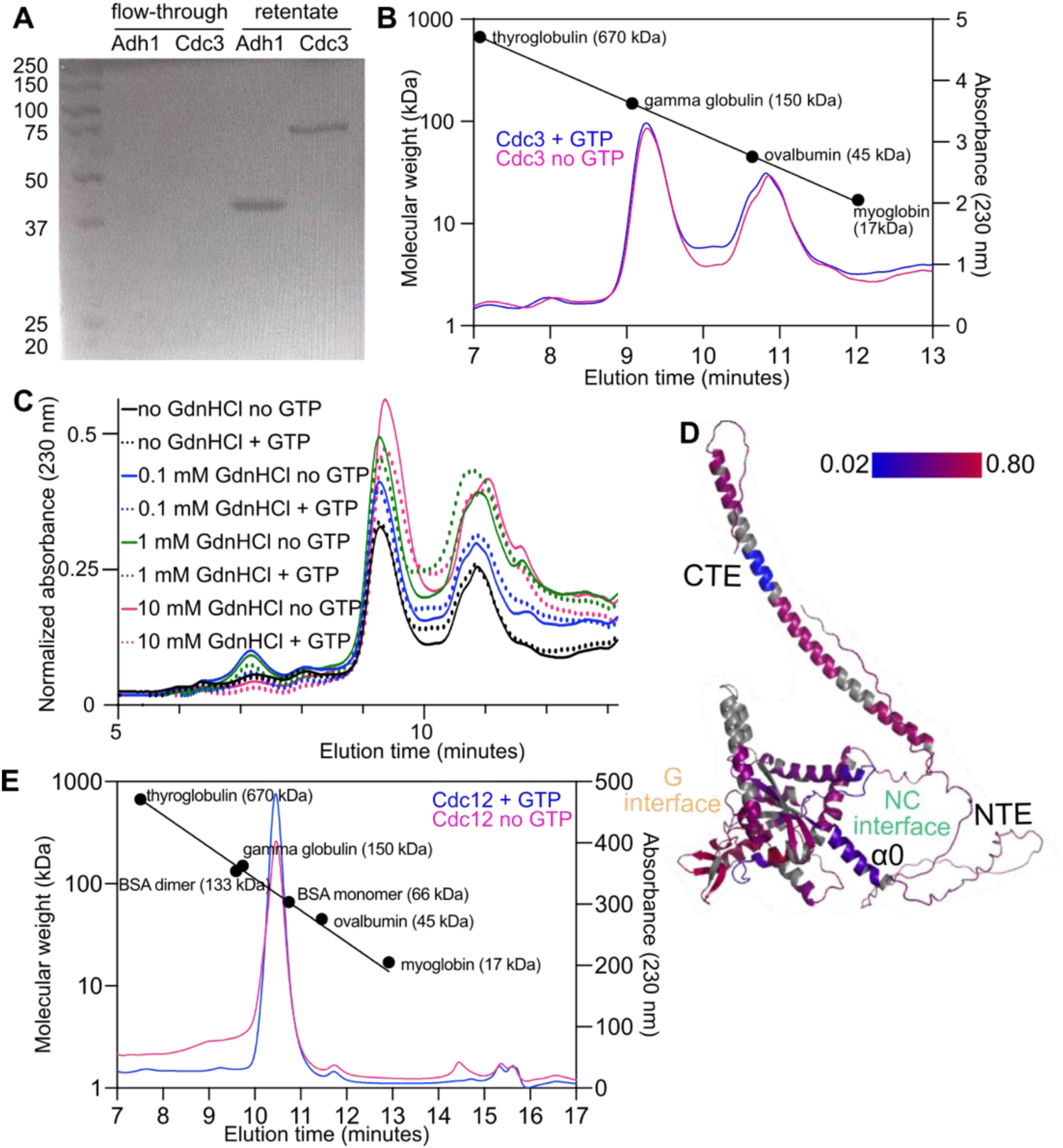
Non-native homodimerization by purified yeast septins. (A) Purified tetrameric Adh1 (native molecular weight ∼150 kDa) and purified 6xHis-Cdc3 (monomeric molecular weight ∼60 kDa) were centrifuged through a 100-kDa-cut-off filtration device. Portions of the flow-through and retentate were separated by SDS-PAGE and stained with Coomassie. Leftmost lane contains molecular weight ladder (Thermo Fisher Scientific # SM0331). (B) Samples of molecular weight standards or 6xHis-Cdc3 were analyzed by size exclusion chromatography and the 230-nm absorbance profiles, indicative of protein concentration, were plotted as a function of elution time. Colored lines show 6xHis-Cdc3 with or without added GTP. Circles show peak elution times for the indicated standards. (C) As in (B), but only with samples of 6xHis-Cdc3 containing or lacking GTP (2 mM) and/or GdnHCl at the indicated concentrations. GdnHCl was also included in the running buffer at the same concentrations. (D) Hydrogen-deuterium exchange ratios after 7200 seconds incubation of 6xHis-Cdc3 in deuterated buffer were mapped onto the predicted structure of Cdc3. Color indicates exchange ratio; regions in gray were not detected by MS. Locations of the CTE, NTE, α0 helix, and G and NC interfaces are for illustration purposes only. (E) As in (B), but with purified Cdc12.

In analytical size exclusion chromatography (SEC), 6xHis-Cdc3 eluted as a ∼1:1 mix of an apparent monomer and homodimer (Figure 1B). The monomer eluted later than 45-kDa ovalbumin (Figure 1B), suggestive of a non-spherical shape and/or interactions with the column media. The apparent dimer eluted as expected (∼120 kDa). These results indicate that in the absence of native interaction partners, Cdc3 is prone to non-native homodimerization. Non-native Cdc3 homodimerization via the “G” interface –– which encompasses the GTP-binding pocket –– allows septin filament formation in the absence of Cdc10 (Johnson et al., 2020; McMurray et al., 2011b). Exogenous addition of GDP or a GTP analog was able to drive G-interface-mediated dimerization of a purified human septin (Sirajuddin et al., 2007) and exogenous guanidine hydrochloride promotes non-native Cdc3 homodimerization *in vivo* (Johnson et al., 2020). We added an excess of GTP and/or guanidine hydrochloride at various concentrations and repeated the SEC analysis. Neither small molecule changed the Cdc3 monomer:dimer ratio (Figure 1C).

To gain further insights into the conformation of Cdc3 and the mechanism of homodimerization, we performed hydrogen-deuterium exchange mass spectrometry (HDX-MS) and mapped exchange ratios onto the Cdc3 amino acid sequence (Supplemental Figure 1) and the predicted structure of Cdc3 (Figure 1D). MS results confirmed that the purified protein was full-length, though some internal peptides were not represented (∼85% coverage; Figure 1D and Supplemental Figure 1). Consistent with the SEC results, every peptide in the HDX-MS data was flagged by the analysis software for likely “EX1” kinetics (poor fit to binomial distribution), pointing to a mixture of at least two distinct conformations.

Among the five septins expressed in mitotically dividing yeast cells, Cdc3 is unique in having a ∼100-residue N-terminal extension (NTE), which exhibits a very high, evolutionarily conserved predicted propensity for intrinsic disorder (Weems and McMurray, 2017). We previously proposed that the Cdc3 NTE interacts weakly with the Cdc3 G interface to inhibit G dimerization with Cdc10 until Cdc3–Cdc12 interaction across the NC interface repositions the NTE and allows Cdc10 access (Weems and McMurray, 2017). Consistent with this model, the entire NTE exhibited high exchange ratios (Figure 1D, Supplemental Figure 1), suggestive of high solvent accessibility. (Note that the position of the NTE in the structural model in Figure 1D is a low-confidence prediction.)

Moving C-terminally along 6xHis-Cdc3, the exchange ratio abruptly decreased near the α0 helix (Figure 1D, Supplemental Figure 1), a key feature of the other septin dimerization interface, called “NC”. Other portions of the NC interface, including the α5 helix, also exhibited slower exchange (Figure 1D, Supplemental Figure 1). We note that the sequences in many regions that exhibited less exchange are rich in hydrophobic amino acids. We interpret these data as evidence that Cdc3 purified in these conditions is at least partially folded, burying hydrophobic/aggregation-prone regions in intra- or inter-molecular contacts, but is mostly in an unstable, largely non-native conformation.

Regions normally buried in the G dimer interface showed relatively high exchange, including the G1 motif/P loop, the Sep4 motif/β7-β8 hairpin, the *trans* loop 1, and the G3 motif/Switch II loop (Supplemental Figure 1). The C-terminal extension, or CTE, displayed high exchange except for a region within the hydrophobic heptad repeats predicted to mediate formation of a coiled-coil (Versele et al., 2004) (Figure 1D, Supplemental Figure 1). This region exhibited the lowest exchange ratios of any part of Cdc3, suggesting stable, efficient burial of these backbone hydrogens. Taken together, these results are most consistent with a model in which, when purified following heterologous expression in the absence of its native interaction partners, the yeast septin Cdc3 forms an unstable non-native homodimer involving the NC interface and a CTE-mediated coiled-coil. These findings agree well with previous data in which interaction of purified 6xHis-Cdc3 with purified GST-Cdc3 required the CTE (Versele et al., 2004). Notably, the only other yeast full-length yeast septin to be previously analyzed at this level of detail, Cdc11, behaved in a similar way, forming a CTE-dependent homodimer (Brausemann et al., 2016). The inability of guanidine or GTP to drive 6xHis-Cdc3 homodimerization across the G interface point to a key role for the presence of native interaction partners (chaperones or other septins) to prevent the acquisition by Cdc3 of non-native conformations prone to non-native interactions.

We examined Cdc12 purified in a similar way, although we had to try a number of approaches before we found one that gave sufficient yield and purity: fusion to an N-terminal maltose-binding protein-7xHis tag that was cleaved off via a TEV protease site. SEC analysis of Cdc12 at ∼2 µM indicated that the majority of Cdc12 eluted as a stable homodimer of ∼92 kDa, independent of added GTP (Figure 1E). A small peak corresponding to apparent molecular weight of ∼35 kDa likely represents a minor fraction of monomeric Cdc12 that, like Cdc3 monomer, eluted “smaller” than its actual size. Thus, Cdc12 is prone to non-native homodimerization to an even greater extent than is Cdc3. Given previous results that interaction between Cdc12-6xHis and GST-Cdc12 involves the Cdc12 CTE, and that the Cdc12 CTE alone is able to weakly homo-oligomerize (Versele et al., 2004), we suspect that non-native homodimerization by purified Cdc12 is also mediated by the NC interface and coiled-coil-forming sequences in the CTE. These data provide direct evidence that, when native interaction partners are unavailable, wild-type yeast septins adopt non-native conformations and engage in non-native interactions.

### *In vivo* site-specific crosslinking of Cdc10 to CCT

If chaperones bind with low affinity to exposed septin oligomerization interfaces, preventing non-native interactions, and are eventually “competed out” by other septins during native septin complex assembly, then there should be specific chaperone binding sites in septin oligomerization interfaces. Our genetic approach using temperature-sensitive mutants pinpointed individual residues in the G interface that, when mutated, prolong chaperone interactions (Denney et al., 2021; Johnson et al., 2015). However, these residues are not necessarily the sites of chaperone binding. Our previous biochemical pull-down (Johnson et al., 2015) and split-YFP (Denney et al., 2021) studies of chaperone–septin interactions also did not directly address where exactly chaperones bind. To learn more about what parts of a septin might require folding assistance from chaperones, we used a site-specific crosslinking approach in living cells.

The genetically encoded photoactivatable amino acid p-benzoyl-l-phenylalanine (Bpa) can replace specific natural amino acids in a protein sequence *in vivo* (Krishnamurthy et al., 2011). We introduced a UAG codon at specific locations in Cdc3 or Cdc10 and induced overexpression of either septin N-terminally tagged with GST and hexahistidine (6xHis) in yeast cells also carrying a plasmid encoding both a modified tRNA that recognizes the UAG codon and an aminoacyl-tRNA synthetase that charges the modified tRNA with Bpa (Krishnamurthy et al., 2011). Sites of Bpa incorporation were chosen based on sequence-based algorithms that predict propensity for aggregation and chaperone binding sites (Figure 2A). Additionally, a non-septin Bpa crosslinking study demonstrated that a β-hairpin is sufficient to recruit the cytosolic Hsp40 chaperone Ydj1 (Jores et al., 2018). The highly conserved septin β7-β8 hairpin, which harbors the “Sep4” motif (Pan et al., 2007), not only overlaps with predicted Hsp70 and Ydj1 binding sites in multiple septins, it is also buried in the septin G interface, and should be exposed until a newly-made septin incorporates into septin hetero-oligomers (Figure 2A).

**Figure 2.**
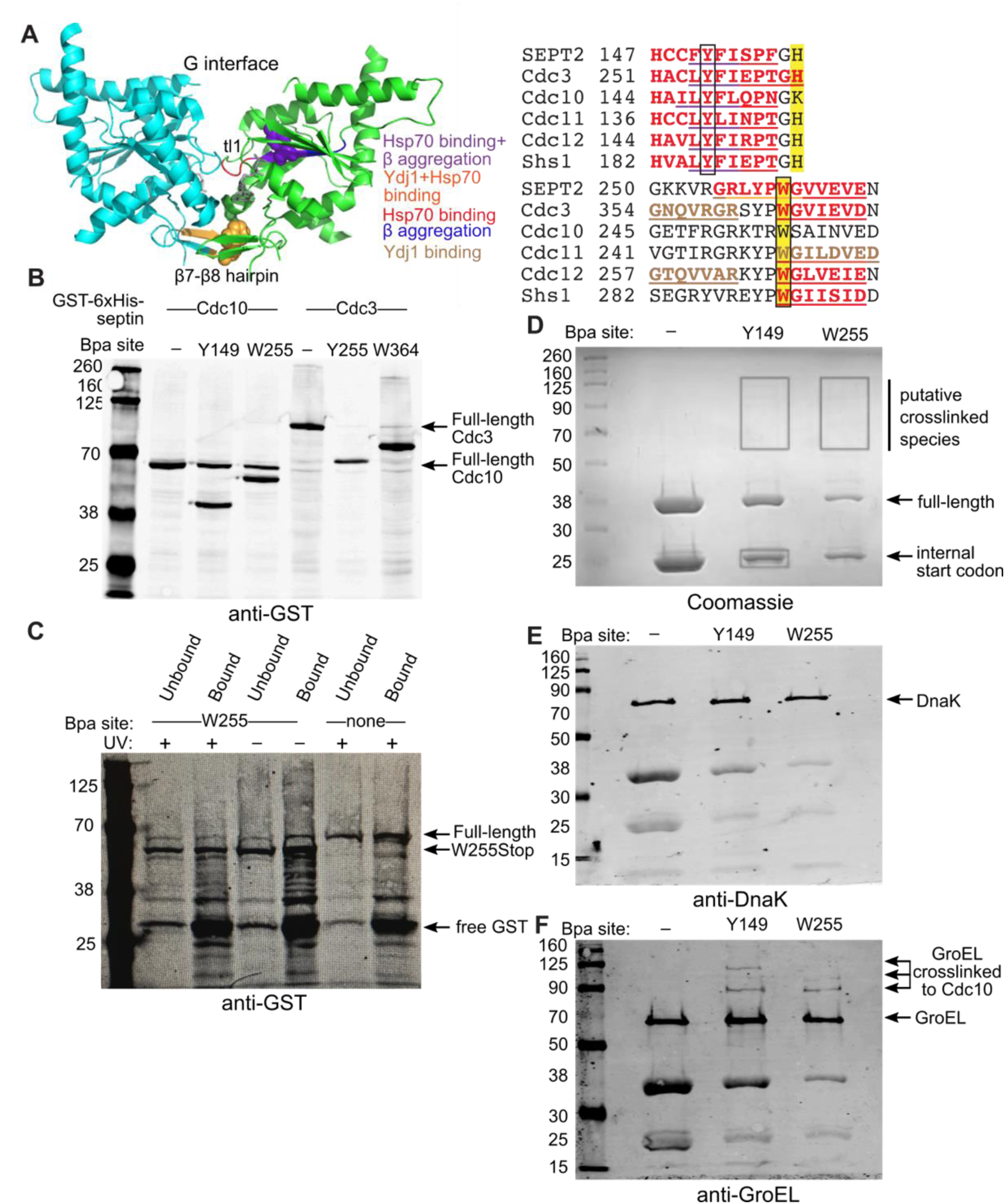
In vivo crosslinking between chaperonins and yeast septins. (A) In the crystal structure of human Sept2 G homodimer (PDB 2QNR), the colored ribbon follows the color scheme in the sequences at right for overlapping β-aggregation, Ydj1 and Hsp70-binding sites, and shows in light orange the β7-β8 hairpin that makes contacts across the septin G interface. “tl1”, *trans* loop 1. Sequence alignments show predicted binding sites underlined and in bold. Predicted Ydj1 binding sites (GX[LMQ]{P}X{P}{CIMPVW} (Kota et al., 2009) are in brown, β-aggregation-prone sequences according to TANGO (Fernandez-Escamilla et al., 2004) are in blue, and where each overlaps an Hsp70 binding site (LIMBO score >11.08 (Van Durme et al., 2009)) is in dark orange or purple, respectively. Highlighted residues make contacts across the G interface. Boxed residues are shown as spheres in the ribbon structure and are the sites chosen for replacement by the photocrosslinker Bpa. (B) Yeast cells (strain YRP2838) carrying plasmid pSNRtRNA-pBPA-RS and a plasmid encoding a GST-6xHis-tagged septin (Cdc3 or Cdc10) with or without sites of incorporation of the photocrosslinker amino acid Bpa were grown in the presence of Bpa and, to induce tagged septin expression, 2% galactose. Total protein was extracted via alkaline lysis and TCA precipitation and separated by SDS-PAGE before immunoblotting with anti-GST antibodies. (C) As in (B) but just for Cdc10 and cells were exposed to UV to induce crosslinking (or not, as indicated). Following protein extraction in denaturing conditions 6xHis-tagged proteins were purified by Ni-NTA affinity. (D) *E. coli* cells (strain BL21(DE3)) carrying plasmid pEVOL-pBpF and a Cdc10-6xHis plasmid with the indicated site of Bpa incorporation were grown in the presence of Bpa. IPTG was added to induce tagged Cdc10 expression, cells were exposed to UV and then lysed in denaturing conditions. Proteins bound to Ni-NTA were separated by SDS-PAGE and stained with Coomassie. Boxes indicate regions of the gel that were excised for mass spectrometry. (E-F) Samples as in (D) were separated by SDS-PAGE then immunoblotted for DnaK (E) or GroEL (F). The leftmost lanes in all gels contained molecular weight ladder (Li-Cor Biosciences # 928-60000).

We chose two aromatic residues for replacement with Bpa: the conserved Trp of the Sep4 “WG” sequence, and a conserved Tyr between the core of the GTPase domain and the *trans* loop 1, which also makes contacts across the G interface (Weirich et al., 2008). If Bpa incorporation fails, translation terminates at the UAG codon. Such premature termination can also destabilize the mRNA, so we used a yeast strain lacking Nam1/Upf1, an RNA helicase required for nonsense-mediated mRNA decay (Wang and Wang, 2008). We monitored by immunoblotting the production of full-length septins as a readout of Bpa incorporation efficiency. Bpa incorporation into Cdc10 was more efficient than for Cdc3 (Figure 2B). For both sites of Bpa incorporation into Cdc10, truncated forms were slightly more abundant than the full-length proteins (Figure 2B).

We UV-treated yeast cells expressing GST-6xHis-Cdc10 with no UAG codon or with a UAG at codon 255. Since Bpa255 should eventually become buried in the G interface with Cdc3, to avoid crosslinking only to Cdc3 and promote crosslinking to chaperones engaging nascent Cdc10, we crosslinked after 1 or 3 hours of induction of expression of the tagged Cdc10. Immobilized metal affinity purification using the 6xHis tag under denaturing conditions followed by SDS-PAGE and immunoblotting with an antibody recognizing the GST tag did not reveal any obvious slower-migrating bands that would have indicated covalently crosslinked species (Figure 2C). In this experiment Bpa incorporation was much less efficient, and very little full-length protein was made. Additionally, the appearance of a ∼30-kDa GST-tagged protein suggested that during the purification steps cellular proteases cleaved in the linker between the GST-6xHis portion (predicted molecular weight 31 kDa) and Cdc10 (Figure 2C).

We therefore turned to mass spectrometry to sensitively detect proteins co-purifying with GST-6xHis-Cdc10(Trp255Bpa). Of a total of 856 proteins identified, 295 (34%) were found in all samples, including the negative controls (two replicates of wild-type GST-6xHis-Cdc10, and one sample with GST-6xHis-Cdc10(Trp255Bpa) but no exposure to UV; Supplemental Table 1). These proteins included Cdc10, as expected; most others are expected to be highly abundant in galactose-grown yeast cells (Pgk1, Gal10, Gal7, ribosomal proteins, etc.). A single protein was absent from all negative controls and present in both timepoints of both independent biological replicates of samples containing GST-6xHis-Cdc10(Trp255Bpa). That protein was the CCT subunit Cct3 (Supplemental Table 1).

### *In vivo* site-specific crosslinking of Cdc10 to GroEL

CCT and GroEL differ in many ways, and GroEL cannot substitute for CCT for the folding of other cytosolic nucleotide-binding cytoskeletal proteins (actin and tubulins (Balchin et al., 2018; Tian et al., 1995a)). Nonetheless, GroEL binds Cdc10 in an *E. coli* lysate-based *in vitro* translation reaction; adding purified CCT displaces Cdc10 from GroEL and creates a CCT–Cdc10 complex (Dekker et al., 2008). Thus an alternative explanation for the lack in those experiments of any CCT effect on Cdc10 conformation in those experiments (as assessed by native PAGE) is that either chaperonin was sufficient to direct Cdc10 folding. To look for evidence that GroEL directly interacts with Cdc10 *in vivo*, we adapted our Bpa crosslinking approach for *E. coli*.

We introduced into BL21(DE3) cells a plasmid encoding Cdc10 with a C-terminal 6xHis tag and a UAG codon at position 149 or 255, and a plasmid encoding the appropriate modified tRNA and aminoacyl-tRNA synthetase. Crosslinking was performed following induction of Cdc10-6xHis expression with IPTG in the presence of Bpa, followed by affinity purification under denaturing conditions and visualization by SDS-PAGE and Coomassie staining (Figure 2D). We expected that the use of a C-terminal affinity tag would guarantee purification of only full-length Cdc10 (37 kDa), but a band of ∼29 kDa was nearly as abundant and was also present in a control sample with no UAG codon (Figure 2D). This band was excised from the gel and subjected to mass spectrometry, the results of which suggested that this product arose from translation initiation at an internal AUG codon corresponding to Met77 in full-length Cdc10 (Supplemental Figure 2). Bands migrating slower than full-length Cdc10-6xHis were faintly visible only in the samples in which Bpa was incorporated into Cdc10 (Figure 2D). Mass spectrometry identified multiple *E. coli* chaperones in the part of the gel that included these bands, including GroEL and the Hsp70 DnaK (Supplemental Table 2). Both chaperones are highly abundant and are common non-specific contaminants of affinity purifications (Bolanos-Garcia and Davies, 2006; Robichon et al., 2011). We analyzed the same samples by immunoblot using anti-GroEL or anti-DnaK antibodies. Only GroEL was present in slowly-migrating, Bpa-dependent bands (Figure 2E,F). Thus both CCT and GroEL directly interact at the same site in the septin Cdc10.

### Slower septin folding/assembly in chaperone-mutant yeast cells

To ask if chaperone function is required for efficient *de novo* septin folding, we used a plasmid in which transcription of C-terminally GFP-tagged Cdc3 is under control of the galactose-inducible *GAL1/10* promoter, and quantified the kinetics of fluorescence accumulation following the addition of galactose to the growth medium (Figure 3A). Localization to rings of septin filaments at yeast bud necks was taken as evidence that Cdc3-GFP folded properly and co-assembled with endogenous untagged septins into native hetero-oligomers. We interpreted fluorescence away from bud necks (in the cytoplasm/nucleus) as a composite of unpolymerized septin hetero-oligomers containing Cdc3-GFP plus “excess” or misfolded Cdc3-GFP monomers. In wild-type cells, very few septin monomers can be found by either biochemical (Frazier et al., 1998) or microscopy-based approaches (Bridges et al., 2014), indicating that septin folding and higher-order assembly are normally very efficient. Accordingly, in our experimental set-up we suspect that transcription and translation are normally the rate-limiting steps for accumulation of fluorescence signal in septin rings. We previously used this same assay to find evidence that a G interface (β7-β8 hairpin) mutation in Cdc3, Gly365Arg, slows folding of the mutant septin to the point that we can detect a delay in the accumulation of septin ring fluorescence over time (Schaefer et al., 2016). Here, we kept Cdc3 wild-type and instead mutated cytosolic chaperones.

**Figure 3.**
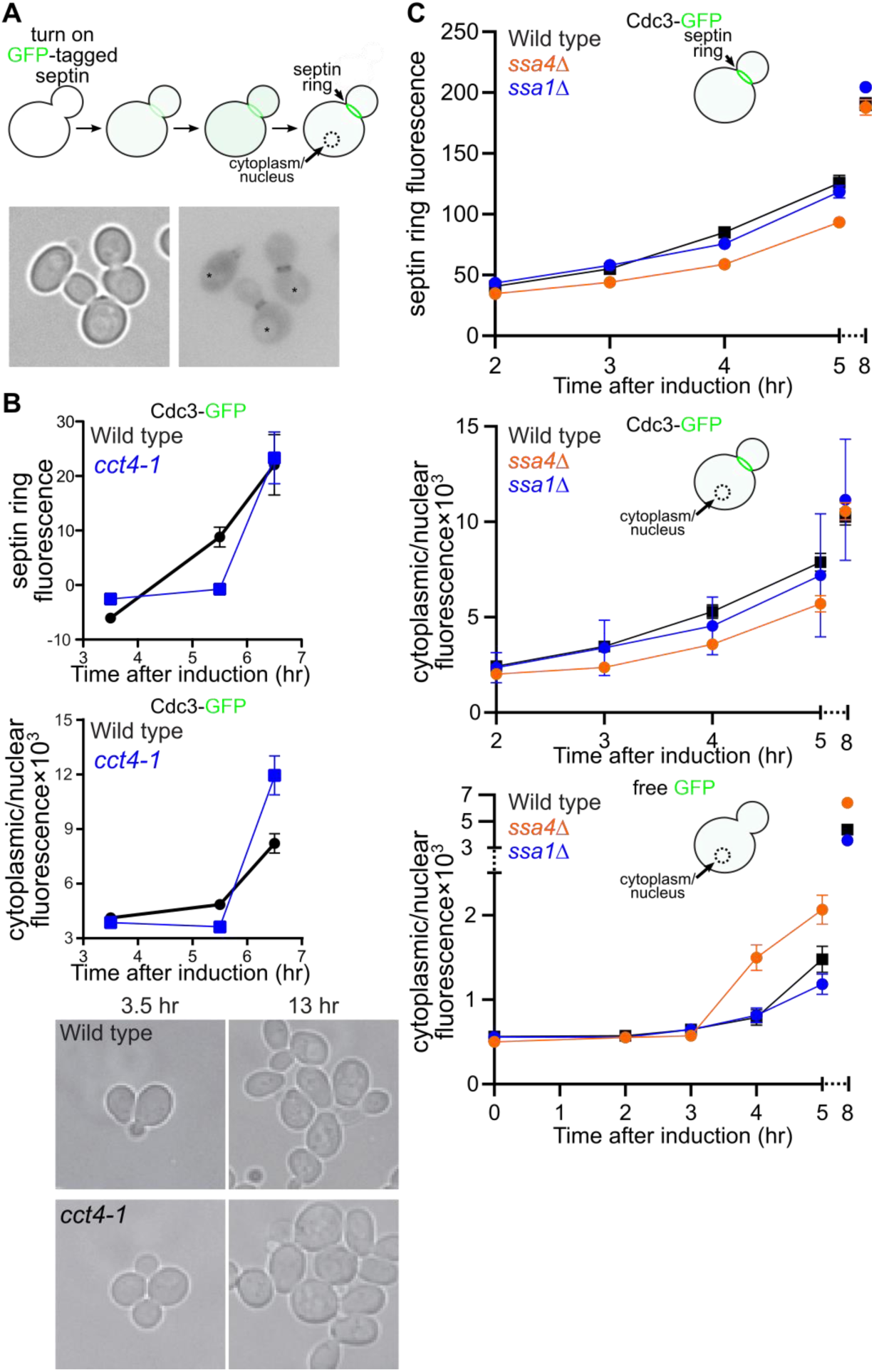
Chaperone requirements for efficient *de novo* septin folding. (A) Schematic illustration of experimental approach. Localization to septin filaments in septin rings at bud necks or in the cytoplasm/nucleus is monitored at timepoints following the induction of new synthesis of a GFP-tagged septin under control of the galactose-inducible *GAL1/10* promoter. Signal is quantified at the bud neck or in a circular region of the cytoplasm/nucleus, avoiding the vacuole. Micrographs below are representative cells from a middle timepoint in an actual experiment. Left panel, transmitted light; right panel, fluorescent signal shown as dark pixels on a light background. Asterisks are centered in presumptive vacuoles. (B) Wild-type cells (BY4741) or *cct4* mutant cells (CBY11211) carrying plasmid pMVB1 were grown in synthetic medium with 2% raffinose and loaded into a microfluidic chamber. Medium containing 1.95% raffinose and 0.05% galactose was introduced and images were taken at the indicated timepoints thereafter. At least 46 cells were analyzed per genotype per timepoint. Points show means, error bars are SEM. The micrographs below show the same fields of cells at two timepoints, demonstrating similar rates of cell division for the wild-type and mutant cells. (C) As in (B) but with strains H00504 (“*ssa1Δ*”) and H00507 (“*ssa4Δ*”) and, as indicated, Cdc3-GFP plasmid pMVB1 or GFP alone plasmid pTS395. Cells were grown in culture tubes and aliquots were removed for imaging at the indicated timepoints. At least 30 cells were analyzed per genotype per timepoint.

We tested CCT using *cct4-1*, in which the Cct4 subunit carries a Gly345Asp substitution that interferes with cooperativity of ATP binding between the two rings of the chaperonin complex (Shimon et al., 2008). Cdc3-GFP mislocalization as misshapen septin rings and aberrant cortical patches was previously seen in the same *cct4* mutant at the restrictive temperature of 37°C, where septins no longer co-purify with the mutant CCT (Dekker et al., 2008). At 37°C the mutant cells also make multiple, elongated buds (Shimon et al., 2008), hallmarks of septin dysfunction. Our assay for septin folding kinetics requires that the mutant cells proliferate at the same rate as wild-type, otherwise differences in fluorescence could be attributed to changes in dilution through cell divisions. Hence we cultured the cells at room temperature (22°C; Figure 3B). We suspected that CCT–septin interactions would be compromised in *cct4-1* cells even at permissive temperatures because: (i) the actin-folding activity *in vitro* of purified CCT carrying mutant Cct4(Gly354Asp) subunits is crippled to the same extent at permissive and restrictive temperatures (Shimon et al., 2008); and (ii) at an otherwise permissive temperature *cct4-1* displays a negative genetic interactions when combined with the slow-folding G-interface mutant septin Cdc10(Asp182Asn) (Costanzo et al., 2016; Johnson et al., 2015). Indeed, the temperature-sensitive nature of the *cct4* mutant may say more about the adverse effects of high temperatures on wild-type protein folding than on Cct4(Gly354Asp) function *per se*.

The accumulation of fluorescent Cdc3-GFP signal in septin rings was delayed in *cct4* mutants relative to wild-type cells, with almost no increase in the mutant cells between the 3.5- and 5.5-hour timepoints post-induction, but by 6.5 hours septin rings were as bright as in wild-type cells (Figure 3B). Previous studies using analogous galactose-induction assays but testing a different client protein demonstrated that CCT function is dispensable for efficient transcriptional induction by the *GAL1/10* promoter (Kim et al., 1998). GFP folds independently of chaperones (Makino et al., 1997; Weissman et al., 1996). Thus these data support a requirement for CCT function in efficient *de novo* septin folding.

Chaperonin assistance is necessary but not sufficient for the *de novo* folding of other cytoskeletal NTPases; other chaperones are also required (Gao et al., 1993; Machida et al., 2021; Vainberg et al., 1998). We previously found interactions between septins and multiple, partly redundant members of the yeast cytosolic Hsp70 chaperone family, including Ssa2, Ssa3, and Ssa4 but not Ssa1 (Denney et al., 2021), which is 98% identical to Ssa2. We tested *ssa1Δ* and *ssa4Δ* mutants and, consistent with our interaction data, found a delay in Cdc3-GFP fluorescence in septin rings in *ssa4Δ* cells but not in *ssa1Δ* (Figure 3C). The delay in Cdc3-GFP accumulation in *ssa4Δ* cells may even have been partially masked by higher induction from the *GAL1/10* promoter at later timepoints, since in these cells free GFP expressed from an otherwise identical plasmid accumulated more quickly than in wild-type or *ssa1Δ* cells (Figure 3C). These data may point to less efficient septin translation in *ssa4Δ* cells. Indeed, inhibition of human Ssa homologs slows translational elongation of many proteins, which has been proposed to reflect loss of a co-translational “pulling force” that these chaperones normally exert on a nascent client peptide emerging from the ribosome exit tunnel (Liu et al., 2013; Shalgi et al., 2013).

The folding activity of Ssa-family chaperones is modified by interaction with either of two cytosolic Hsp40-family chaperones, Ydj1 or Sis1 (Lu and Cyr, 1998), both of which bind wild-type septins (Denney et al., 2021). Sis1 is essential (Luke et al., 1991) and *ydj1Δ* cells divide slowly (Caplan and Douglas, 1991) so we tried to query the involvement of these chaperones in Ssa function on septin folding kinetics using point-mutant Hsp40– Hsp70 pairs that “force” a single Ssa chaperone to interact only with a specific Hsp40 (Reidy et al., 2014) (Supplemental Figure 3A). However, a Ssa2(Arg169His)– Ydj1(Asp36Asn) combination increased *GAL1/10* promoter activity, as evidenced by higher levels of both Cdc3-GFP and free GFP (Supplemental Figure 3A), precluding our analysis. Finally, we found no difference in Cdc3-GFP folding kinetics in cells lacking either of the two partially redundant cytosolic Hsp90 family members, Hsc82 and Hsp82 (Supplemental Figure 3B). Since the range of effects detectable by our kinetic assay is rather narrow –– chaperone mutations must not be so severe as to slow cell division or perturb septin ring morphology but severe enough to cause a phenotype –– the failure to detect an effect cannot be interpreted as a lack of chaperone involvement in septin folding.

### Chaperone requirements for post-translational septin incorporation into native hetero-oligomers

The absence of native interaction partners predisposes septins to non-native conformations and interactions. Are these septins able to make native interactions if they encounter native partners long after they were synthesized? If cytosolic chaperones co-translationally promote native septin conformations, then individual septins synthesized in the cytosol at great excess compared to the levels of other septins may also adopt non-native conformations and have difficulty interacting properly with other septins once those native partners become available. In other words, the excess septin may become “distracted” while it waits. Chaperones could also act post-translationally to protect an excess septin from non-native conformations and maintain the ability to subsequently interact with native partners long after synthesis. We used the *GAL1/10* promoter to individually overexpress single fluorescently-tagged septins and monitored their ability to incorporate into septin filaments after blocking new expression by the addition of glucose (Figure 4A). Excess GFP- or mCherry-tagged molecules of Cdc3, Cdc12, or the slow-folding mutant Cdc3(G365R) accumulated in the cytoplasm/nucleus and, as cells divided without new expression of the tagged septin, cytoplasmic/nuclear signal decreased approximately exponentially with each cell division (Figure 4B-E), consistent with dilution and negligible degradation (McMurray and Thorner, 2008). By contrast, septin ring fluorescence decreased more slowly than the cytoplasmic/nuclear signal (Figure 4B-E), indicating that, many hours after they were translated, the excess tagged septin proteins were able to assemble into complexes capable of polymerizing into septin filaments

**Figure 4.**
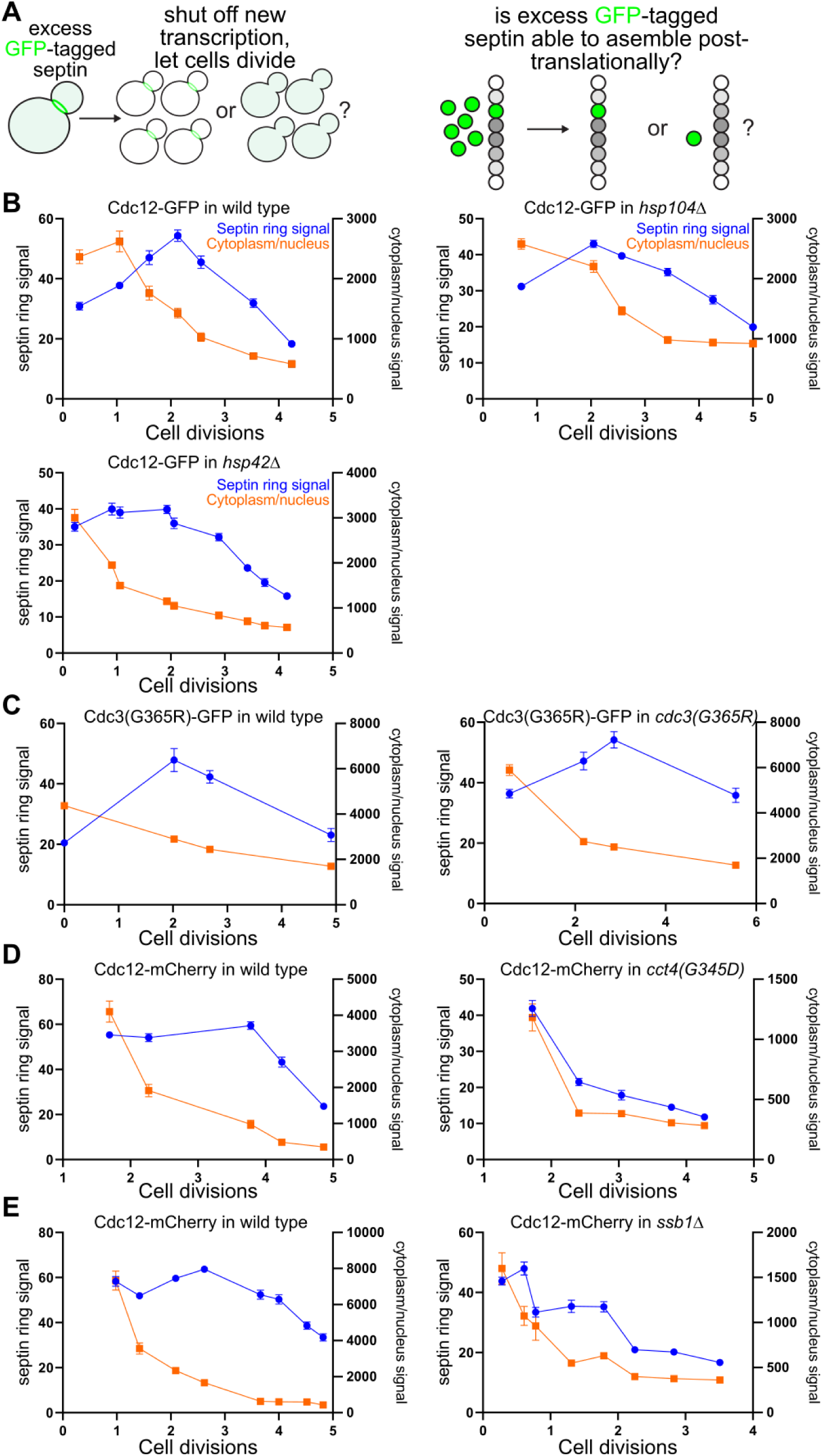
Chaperone requirements for post-translational septin assembly. (A) Schematic illustration of experimental approach. An excess of a single, fluorescently tagged septin is generated by transient overexpression from the galactose-inducible, glucose-repressible *GAL1/10* promoter, and the fluorescent signal in septin filaments at the bud neck or in the cytoplasm/nucleus is quantified via microscopy at various timepoints after cessation of new expression (addition of glucose). If the excess septin is capable of assembling with newly made partner septins long after it was synthesized, then the bud neck signal should increase initially and decrease slowly thereafter via a mix of incorporation into septin filaments and dilution through cell divisions. If, on the other hand, the excess septin is unable to assemble with other septins, bud neck signal should decrease exclusively via dilution, similar to the more rapid and approximately exponential kinetics expected for the cytoplasmic/nuclear signal. (B) Wild-type diploid cells (BY4743), *hsp104Δ/hsp104Δ* mutant cells (strain 31514), or *hsp42Δhsp42Δ* mutant cells (diploid made by mating strains H00481 and H00419) carrying plasmid pMVB2 were grown in synthetic medium with 0.1% galactose and 1.9% raffinose. Following the addition of glucose to final 2%, aliquots were taken at timepoints and imaged (50 cells per genotype per timepoint). Cell concentration in the culture was also monitored at each timepoint to calculate the number of cell divisions following glucose addition. Points show means, error bars are SEM. (C) As in (B) but with haploid strain BY4741 (“wild type)” or CBY07236 (“*cdc3(G365R)*”) carrying plasmid pPgal-Cdc3(G365R)-GFP, and with 40 cells per genotype per timepoint. (D) As in (C) but with plasmid pGF-IVL-470 and *cct4* strain CBY11211. At least 33 cells were imaged per genotype per timepoint. (E) As in (B) but with plasmid pGF-IVL-470 and *ssb1Δ* mutant haploid cells (strain H00421). 40 cells per genotype per timepoint.

We noticed fluorescent foci in a subset of cells overexpressing Cdc12-GFP (Supplemental Figure 4A,B) and suspected they could be aggregates of misfolded septin. However, further experiments suggested that these foci reflect properly-folded Cdc12 artificially oligomerized by the GFP tag. First, the foci frequently recruited other septins, but rarely recruited the disaggregase Hsp104 (Supplemental Figure 4A). By contrast, Hsp104 co-localized with most foci formed by overexpressing the misfolding-prone mutant septin Cdc10(D182N) (Supplemental Figure 4A), which we have previously shown to interact strongly with Hsp104 (Denney et al., 2021; Johnson et al., 2015). Deleting *HSP104* slightly increased Cdc12-GFP focus formation but foci disappeared just as quickly in the absence of Hsp104 as in its presence (Supplemental Figure 4B), arguing against the idea that the foci are substrates of Hsp104 disaggregase activity. Second, while N-terminally GST-5xHis-tagged Cdc12 was also able to induce the formation of foci and long “rods” by endogenous Cdc10-mCherry, overexpressed Cdc12-mCherry did not form foci or rods (Supplemental Figure 4C).

To ask if chaperones are required for efficient septin post-translational hetero-oligomerization, we transiently overexpressed tagged Cdc12 in chaperone-mutant cells and monitored the kinetics of signal decay. Deleting *HSP104* did not affect the ability of excess Cdc12-GFP to incorporate into septin filaments (Figure 4B). In *cct4* cells, on the other hand, we found a clear defect in post-translational Cdc12 incorporation, with both septin ring and cytoplasmic/nuclear signals decreasing approximately exponentially with each cell division, consistent with dilution (Figure 4D). Together with the apparent delay in *de novo* folding we observed in *cct4* mutants (Figure 3B), we interpret these results to mean that when CCT is even slightly dysfunctional, newly-made septins are: (i) slower to achieve the conformation(s) required for efficient septin-septin interactions, and (ii) less able to achieve and/or maintain quasi-native conformations when septin-septin interactions are further delayed by a scarcity of available septin partners, as in the case of individual septin overexpression.

Two paralogous ribosome-associated Hsp70 chaperones, Ssb1 and Ssb2, interact co-translationally with over half of all actively translated proteins, including Cdc3 and Cdc12, and all septins are found in aggregates in *ssb1Δ ssb2Δ* cells (Willmund et al., 2013). The enrichment of subunits of oligomeric complexes among nascent Ssb-interacting polypeptides was suggested to reflect “a potential role […] in stabilizing free subunits of oligomeric complexes, which may display a higher number of exposed contact interfaces” (Willmund et al., 2013). While Ssb1 and Ssb2 are usually considered functionally redundant, we noticed in *ssb1Δ SSB2*^+^ cells an increase in apparent aggregates (intracellular foci) when we overexpressed the human-disease-linked mutant Cdc10(D182N)-GFP and cultured cells at 37°C (Supplemental Figure 4D), pointing to a functional defect in the absence of Ssb1 alone. As with *cct4* mutants, in *ssb1Δ* cells excess Cdc12-mCherry molecules were unable to efficiently incorporate into septin filaments long after their synthesis (Figure 4E). We noticed “leaky” expression of Cdc12-mCherry in glucose cultures of *ssb1Δ* cells (Supplemental Figure 4E), which is consistent with known glucose repression defects in *ssb* cells (Dombek et al., 2004; Hübscher et al., 2016; von Plehwe et al., 2009) and likely explains why after glucose addition Cdc12-mCherry signal in septin rings did not decrease as rapidly as it did in *cct* cells. Thus our data likely underestimate the *ssb* septin defect. We also tested deletion of *HSP42*, encoding the general small heat shock protein of the yeast cytosol (Haslbeck et al., 2004). We had noticed that loss of Hsp42 eliminated foci of overexpressed Cdc10(D182N)-GFP (Supplemental Figure 4D). The absence of Hsp42 had no effect on post-translational Cdc12 assembly (Figure 4B). Taken together, these results point to CCT and Ssb as chaperones that engage recently translated septins and promote conformations capable of interacting properly with partner septins long after synthesis.

### GroEL dysfunction inhibits heterologous production of the yeast septin Cdc12

Our data with chaperone-mutant yeast cells pointed to requirements for CCT and cytosolic Hsp70-family chaperones in efficient septin folding, but since the pathway of septin complex localization to the bud neck is not completely understood, it was possible that the chaperone mutations compromised the function of some other protein(s) that normally promote septin localization. As an independent approach, we mutated cytosolic chaperones in *E. coli* and looked for defects in hetero-oligomerization by heterologously expressed yeast septins. Based on our yeast septin-chaperone interaction results (Denney et al., 2021; Johnson et al., 2015), we focused on ClpB (homolog of the disaggregase Hsp104), DnaJ (major cytosolic Hsp40), DnaK (sole cytosolic Hsp70), GroEL, and HtpG (sole cytosolic Hsp90).

For *ΔclpB* and *ΔhtpG* we obtained *E. coli* deletion strains (Baba et al., 2006) representing these non-essential chaperones and lysogenized them with the DE3 prophage to allow T7 RNA polymerase expression upon addition of IPTG. Since DnaJ and DnaK are required for efficient phage replication (Yochem et al., 1978), we first lysogenized the wild-type parent strain of the *E. coli* deletion collection and then introduced *ΔdnaJ* or *ΔdnaK* alleles by recombination. In all four deletion strains we took extra steps to rule out the acquisition of extragenic suppressors or the persistence of wild-type alleles (see Methods). We then introduced two plasmids each encoding two yeast septins (Cdc10 and 6xHis-tagged Cdc12 or Cdc3 and Cdc11) under control of the T7 promoter. For GroEL, we used a *ΔgroEL* strain carrying three plasmids: one encoding the temperature-sensitive *groEL(Glu461Lys)* mutant allele (Chapman et al., 2006), one encoding T7 RNA polymerase under control of the IPTG-inducible *lac* promoter, and one encoding untagged Cdc3 and 6xHis-tagged Cdc12 under T7 promoter control. Cells were grown at 24°C to mid-log phase and subsequently induced with IPTG at 24°C or 37°C. Following cell lysis, 6xHis-Cdc12 was purified from the soluble fraction and stoichiometric co-purification of the other, untagged septins was taken as evidence of proper septin folding and complex assembly. Stoichiometric septin complexes were obtained from *ΔclpB*, *ΔdnaJ*, *ΔdnaK*, and *ΔhtpG* cells (Figure 5A), indicating that these chaperones are not strictly required for *de novo* septin folding, at least in this context. Reductions in the total yield of septin complexes in several mutants (Figure 5A) likely reflect defects in transcriptional induction, rather than in septin translation or folding, since similar reductions were seen for expression of a control protein (see Methods).

**Figure 5.**
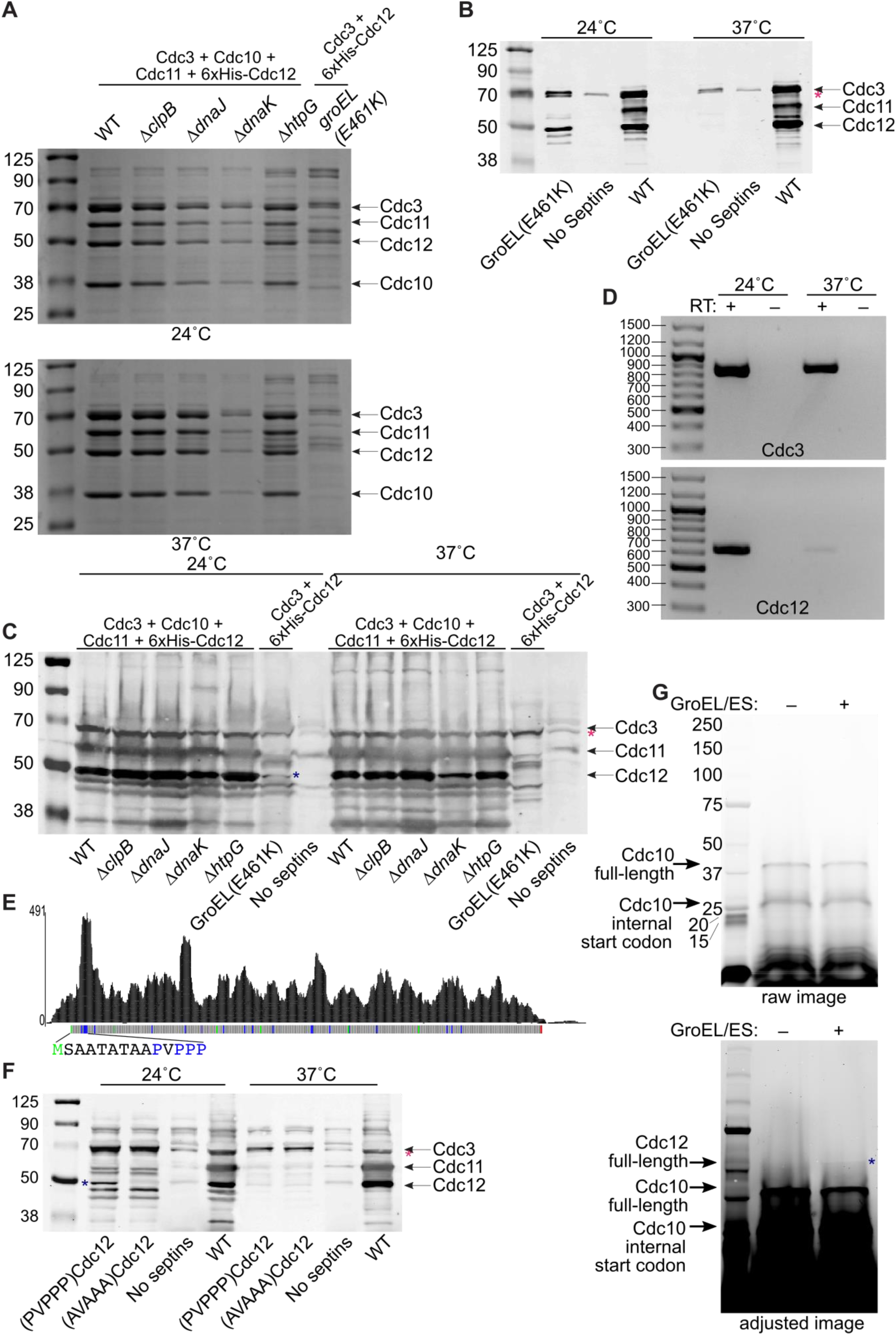
Chaperonin requirement for translation of the septin Cdc12 by prokaryotic ribosomes. (A) Coomassie-stained SDS-PAGE of proteins purified via Ni-NTA from *E. coli* cells carrying plasmids encoding the indicated septins, plus other plasmids as necessary for induction or viability. Strains were B002 (“WT”), B003 (“*ΔclpB*”), B004 (“*ΔdnaJ*”), B005 (“*ΔdnaK*”), B006 (“*ΔhtpG*”), and 461 (“*groEL(E461K)*”). Septin-encoding plasmids were pMVB128, pMVB133, and pMAM54. Leftmost lane contains a molecular weight ladder (Li-Cor Biosciences # 928-60000). (B) Immunoblot of bound proteins prepared as in (A), plus a sample from a strain carrying no septi-encoding plasmid (“No Septins”), using a cocktail of antibodies recognizing Cdc3, Cdc11, and the 6xHis tag. Magenta asterisk indicates a ∼70 kDa protein weakly reactive with the Cdc3 and/or Cdc11 antibody. (C) As in (B) but for urea-solubilized pellets (cell debris and insoluble material) obtained following cell lysis. The blue asterisk indicates the band corresponding to Cdc12. (D) DNA agarose gel electrophoresis and ethidium bromide staining of products of RT-PCR for Cdc3 or Cdc12 mRNA from total RNA extracted from strain 461 carrying plasmid pMAM54 and induced with IPTG at the indicated temperatures. The presence or absence of reverse transcriptase (“RT”) in the reactions is indicated. Leftmost lane contains molecular weight ladder (Thermo Fisher Scientific # SM0331). (E) Published Ribo-seq/ribosome profiling data (Sen et al., 2015) showing the positions of ribosomes along the Cdc12 mRNA as numbers of reads of ribosome-protected mRNA fragments. Below, the Cdc12 ORF is illustrated with Met codons in green, Pro codons in blue, and the stop codon in red. The first 14 amino acids of Cdc12 are indicated with the same color scheme. (F) As in (B) but including samples from *groEL*-mutant cells carrying the plasmid pMAM88, which encodes Cdc3 and 6xHis-tagged Cdc12 with the four Pro residues highlighted in (E) mutated to Ala (“(AVAAA)Cdc12”). Samples from cells with pMAM54, encoding wild-type Cdc12, are indicated by “(PVPPP)Cdc12”. The blue asterisk indicates the band corresponding to Cdc12. (G) Otherwise chaperone-free *in vitro* transcription/translation reactions lacking release factors (PURExpress ΔRF123) and containing fluorescent puromycin to label translation products were resolved by SDS-PAGE. The template plasmid pMVB128 encodes Cdc10 and 6xHis-Cdc12. Purified GroEL/ES chaperonin was added to one reaction, as indicated. Brightness and contrast in the “adjusted image” below were adjusted in FIJI. The blue asterisk indicates the band corresponding to full-length Cdc12. Leftmost lane contains a molecular weight ladder (Bio-Rad #1610395).

By contrast, in the *groEL*-mutant cells bands at the positions expected for Cdc3 and 6xHis-Cdc12 were visible in Coomassie-stained samples from cells induced at 24°C but not at 37°C (Figure 5A). Since we saw multiple other proteins of unexpected sizes in the bead-bound samples, to improve our detection of septin expression we used immunoblotting and antibodies recognizing Cdc3 or the 6xHis tag. These experiments confirmed that, while Cdc3 was present in cells induced at either temperature, 6xHis-Cdc12 was only found in cells induced at 24°C (Figure 5B). Since Cdc12 was also missing from urea-solubilized pellets (Figure 5C), its absence from bead-bound samples was not the result of Cdc12 aggregation/insolubility. These findings suggested that severe GroEL dysfunction perturbs Cdc12 translation. Poor translation of non-*E. coli* proteins heterologously expressed in *E. coli* is frequently accompanied by loss of the mRNA (Boël et al., 2016). Indeed, we found that whereas Cdc3 mRNA was present at similar levels in *groEL*-mutant cells induced at either temperature, 6xHis-Cdc12 mRNA was barely detectable at 37°C (Figure 5D). These results point to a specific defect in Cdc12 translation upon GroEL dysfunction leading to turnover of the Cdc12 mRNA, consistent with a co-translational requirement for GroEL in Cdc12 folding.

### A cluster of prolines near the N terminus promote Cdc12 translation in *groEL*-mutant *E. coli*

Defects in co-translational folding in *E. coli* due to chaperone mutations are known to cause premature termination of translation of some endogenous proteins via recruitment of the ribosome release factor RF3 (Zhao et al., 2021), but it was not obvious why Cdc12 would be affected more than Cdc3. mRNA degradation is also not known to follow RF3 recruitment. Translation problems and loss of mRNA during *E. coli* expression of non-septin proteins correlates with codon content (Boël et al., 2016). Moreover, obligate GroEL clients appear to accumulate “suboptimal” codons over evolutionary time, suggesting that assistance from GroEL is able to overcome otherwise deleterious effects of codon content on translation and folding (Warnecke and Hurst, 2010). If Cdc12 is enriched for “suboptimal” codons compared to Cdc3, this might explain the more severe effect of GroEL dysfunction on Cdc12. However, we found no obvious differences in the predicted “codon adaptation index” of Cdc3 versus Cdc12 (Supplemental Figure 5A). Cdc12 has none of 17 pairs of adjacent codons that reduce translation (Gamble et al., 2016). mRNA secondary structure within the ORF also correlates with slower translation elongation (Burkhardt et al., 2017) but predictions of ORF mRNA secondary structures for Cdc12 and Cdc3 did not reveal any clear differences (Supplemental Figure 5B). We looked for specific pairs of amino acids (Ahmed et al., 2020) known to induce translational pausing in yeast, but again found no clear difference between the two septins (Supplemental Figure 5C). Positively-charged residues, especially Arg, and overall charge, as indicated by isoelectric point, correlate with slow translation of yeast proteins (Riba et al., 2019), presumably due to electrostatic interactions with the negatively-charged ribosome exit tunnel (Charneski and Hurst, 2013; Lu and Deutsch, 2008). Cdc12 has nearly twice the number of Arg residues (29) than does Cdc3 (17), despite being ∼30% shorter (Supplemental Figure 5C), and has a much higher isoelectric point (8.23 vs 5.14). Thus an enrichment of positively-charged amino acids in Cdc12 may explain its susceptibility to translation failure when GroEL is dysfunctional.

We also considered Pro residues, which are intrinsically difficult to translate (Pavlov et al., 2009), and noticed that while overall proline content is similar between Cdc3 (18 Pro residues, 3.5%), Cdc10 (9 Pro residues, 3%), Cdc11 (13 Pro residues, 3%) and Cdc12 (20 Pro residues, 5%), Cdc12 is unique in having a Pro-rich cluster (ProValProProPro), located near the N terminus (Figure 5E). Examination of published yeast ribosome profiling data (Sen et al., 2015) revealed that the major peak of ribosome accumulation on Cdc12 mRNA is at the Pro-rich cluster (Figure 5E). To ask what contribution the Pro-rich cluster makes to the apparent translational defect we observed for Cdc12 in *groEL*-mutant *E. coli*, we mutated all of the Pro residues in the cluster to Ala. If the Pro-rich cluster drives ribosome stalling to the point of translation failure and mRNA degradation, then the mutant construct should restore Cdc12 synthesis. Instead, eliminating the Pro-rich cluster exacerbated the Cdc12 synthesis defect: no mutant Cdc12 was detectable at 24°C, a temperature permissive for culture growth and for synthesis of Cdc3 and wild-type Cdc12 (Figure 5F). The Pro-mutant Cdc12 was translated efficiently and folded normally in wild-type *E. coli*, as evidenced by co-purification with co-expressed Cdc3 (Supplemental Figure 6). Note that the Glu461Lys mutation perturbs GroEL function to some extent at all temperatures, since mutant cells are unable to proliferate in minimal medium at any temperature (Horwich et al., 1993). Whereas for wild-type Cdc12 GroEL dysfunction is only manifested as a synthesis defect at 37°C, even slight dysfunction was enough to block synthesis of the Pro-mutant Cdc12. Thus, rather than conferring sensitivity to translational inhibition accompanied by mRNA degradation, the Pro-rich cluster in Cdc12 confers protection.

### GroEL/ES is sufficient to promote Cdc12 translation during cell- and chaperone-free *in vitro* translation

To ask if the lack of Cdc12 protein synthesis in *groEL*-mutant cells reflects intrinsic difficulties in translating Cdc12 protein without functional GroEL, we attempted to synthesize Cdc12 in the context of a cell- and chaperone-free *in vitro* translation system based on components purified from *E. coli* (Shimizu et al., 2001). The PURExpress system has been used previously to demonstrate co-translational action of GroEL on an established client protein (Ying et al., 2006). To specifically detect translation products, we used a version of this system lacking ribosome release factors and added fluorescently-labeled puromycin at a low concentration, which results in covalent C-terminal fluorescent labeling of translation products (Kawahashi et al., 2007). Cdc12 and Cdc10 mRNAs were generated in the same reactions using T7 RNA Polymerase and a plasmid encoding both Cdc12 and Cdc10, each under control of a T7 promoter. We saw fluorescent bands corresponding to the size of full-length Cdc10 (∼37 kDa) and the short version of Cdc10 made from the internal start codon (∼29 kDa) but nothing the size of full-length Cdc12 (∼49 kDa) (Figure 5G), providing an independent demonstration that Cdc12 is more sensitive than other yeast septins to the absence of GroEL. Note that the pI of Cdc10, 5.47, is similar to that of Cdc3.

To ask if GroEL folding activity is sufficient to restore Cdc12 translation, we obtained purified GroEL and GroES (a co-chaperonin protein required for full GroEL folding activity (Goloubinoff et al., 1989a, 1989b)) from a commercial source, verified *in vitro* ATPase activity and folding activity of the reconstituted holoenzyme on a known client (Supplemental Figure 7), and added it to the septin synthesis reactions. Upon the addition of GroEL/ES, we saw a faint fluorescent band the size of full-length Cdc12 (Figure 5G). Using mass spectrometry, we detected peptides at or near the C termini of Cdc10 and Cdc12 in a GroEL/ES-supplemented reaction (Supplemental Figure 8A, Supplemental Table 3), confirming the production of full-length Cdc10 and Cdc12. Notably, we also found a few peptides from DnaK, presumably a contaminant of the purified GroEL and/or GroES (Supplemental Figure 8A, Supplemental Table 3). We further note that because RF3 is among the release factors missing from these reactions, RF3 recruitment to ribosomes translating Cdc12 cannot explain the defect in Cdc12 synthesis in the absence of GroEL.

The conserved bacterial elongation factor P (EF-P) is required for efficient translation of proteins containing Pro-rich clusters *in vivo* and in the same kinds of cell-free *in vitro* reactions that we used (Ude et al., 2013). To determine to what extent the absence of EF-P from the reactions contributed to defects in Cdc12 translation, we obtained EF-P from a commercial source, added it to reactions, and saw no effect (Supplemental Figure 8B). The lack of a detectable effect of EF-P on Cdc12 was not entirely unexpected, considering that for a different protein with a very similar Pro-rich cluster (ProProProIlePro), an EF-P effect was only apparent at very short reaction times (Ude et al., 2013). Only ∼85 endogenous *E. coli* proteins strictly require GroEL for *de novo* folding; for most GroEL clients, other chaperones like trigger factor and/or DnaK can compensate to some extent for the loss of GroEL (Kerner et al., 2005), and *vice versa* (Zhao et al., 2021). We suspect that the lack of these other chaperones from the *in vitro* reactions explains why adding GroEL/ES was only able to support a low yield of Cdc12. Taken together with other findings, these observations provide direct, independent support for a model in which all septins benefit from co-translational folding assistance from a subset of cytosolic chaperones, including a chaperonin, with Cdc12 being especially sensitive to defects in chaperonin function.

## DISCUSSION

It is easy to imagine that protein quaternary structure arises directly and spontaneously from proper tertiary structure, but reality is more complicated, especially in living cells. Since native tertiary structure is often only achieved in the context of quaternary structure, the two are not easily resolved. We focus on septin proteins, which are both more complex and less studied than other, simpler cytoskeletal proteins like the tubulin hetero-dimer. More subunits per complex (up to eight) and a propensity for homo-oligomerization means that there are many “wrong” ways to assemble septin subunits, such as what happened in early *in vitro* studies of purified budding yeast septins (Versele et al., 2004). In the end, the purified complexes that revealed the true subunit organization within septin complexes were made either by co-expression of all the subunits in the same cells (Bertin et al., 2008), or, tellingly, by expression of different subunits in different cells and mixing the cells together during cell lysis (Mendonça et al., 2019, 2021). The failure to achieve native assembly when mixing purified septins compared with the success when co-expressing septins or mixing cells points to some cellular factor(s) important for native septin assembly. Our findings provide evidence that those factors are cytosolic chaperones.

Our observations of apparent septin folding defects in *cct* and *ssb* mutants fit well with proposed functions of CCT and Ssb in co-translationally engaging free subunits of multiprotein assemblies. A class of non-septin CCT clients enriched in subunits of protein complexes bind relatively slowly to the chaperonin during their translation, which was suggested to reflect “the action of cooperating upstream chaperones delaying binding to [CCT] during translation” and client “accumulation” on CCT prior to “release into oligomeric assemblies” (Yam et al., 2008). Given that Ssb binds co-translationally to nascent septins upstream of where CCT binds (Stein et al., 2019), the simplest explanation of our results is that nascent septins are first engaged by Ssb, presumably in complex with its J-protein partner, Zuotin/Zuo1 (Yan et al., 1998), and the “atypical” Hsp70 Ssz1 (Lee et al., 2021), which does not bind septins (Denney et al., 2021) (Figure 6). Early septin folding steps bury the sequences that recruited Ssb, triggering Ssb release and allowing engagement by CCT, which recognizes a septin conformation nearer to the native state. ATP hydrolysis by CCT releases a quasi-native septin from the chaperonin chamber and, if a partner septin is available for interaction, the partner septin occupies the site of CCT binding and prevents further rounds of CCT engagement (Figure 6). If no partner septin is available, rounds of CCT binding and release continue until one is. CCT dysfunction inhibits post-translational septin assembly because the “free” septin falls into kinetic traps – non-native, low-free-energy conformations that are not competent for septin-septin interaction and are susceptible to prolonged association with other chaperones as well as non-native intramolecular interactions. Ssb dysfunction leaves nascent septins in non-native conformations that are aggregation-prone and unrecognizable to CCT.

**Figure 6.**
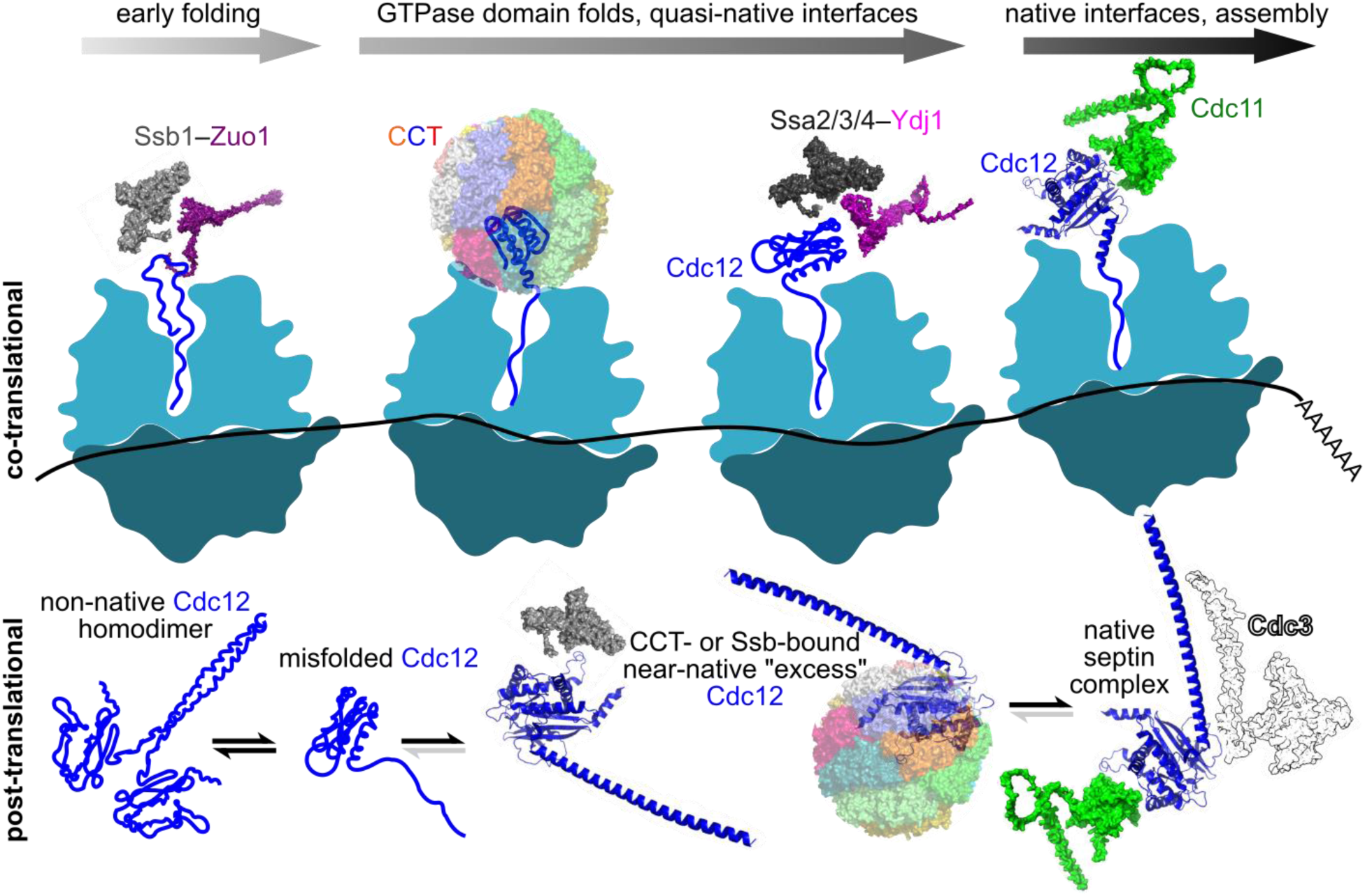
Model for co-translational chaperone-mediated septin folding in the context of septin complex assembly. In this illustration, folding and translation proceed from left to right. CCT and Ssa-family chaperones (but not Ssa1) interact with nascent Cdc12 to promote native folding until Cdc12 has achieved a conformation capable of stably interacting with partner septins. Ydj1 and Zuo1 are shown as presumptive Hsp40s. If folded Cdc11 is available during Cdc12 translation, a Cdc12–Cdc11 heterodimer assembles co-translationally (“co-translational”). Alternatively, if Cdc12 is in excess to other septins, it completes translation without having interacted with another septin (“post-translational”). Here, Cdc12 is susceptible to “off-pathway” non-native conformations prone to non-native homodimerization. CCT and Ssb either continue to interact with full-length Cdc12 (as illustrated), inhibiting off-pathway conformations until stable septin complexes assemble, or (not shown) their dysfunction during translation produces full-length Cdc12 that is irreversibly misfolded and has no path to stable septin assembly. Other septins presumably require similar chaperone assistance for *de novo* folding but Cdc12 may be the slowest to be translated and thus a potential “platform” for co-translational assembly. Protein structures are AlphaFold predictions (the NTE of Cdc3 is hidden, for clarity) or a cryo-EM structure of human CCT (PDB 7LUM) (Knowlton et al., 2021).

The chaperone prefoldin acts similarly for actin folding: the absence of prefoldin during actin translation leads to an actin folding state that is not efficiently folded by CCT, even when prefoldin is added post-translationally (Machida et al., 2021). While we previously found prefoldin binding to septins (Denney et al., 2021) and saw effects of prefoldin mutations on mutant septin oligomerization (Johnson et al., 2015), we did not examine prefoldin here because *E. coli* lacks an obvious prefoldin equivalent, hence septin folding does not strictly require prefoldin assistance. We note that whereas the failure of a “free” subunit to fold efficiently and bind stably to assembly partners targets many other kinds of proteins for proteolytic degradation (Goldberg, 2003; Goldberg and Dice, 1974), yeast septins somehow evade this fate.

Generally speaking, Ssa chaperones interact with nascent chains but to a lesser extent than do Ssb and CCT (Albanèse et al., 2006), so Ssa4 may act downstream of CCT to promote efficient septin folding (Figure 6). The folding delay in *ssa4Δ* but not *ssa1Δ* cells is consistent with our physical septin-chaperone interaction assays (Denney et al., 2021) and, since under non-stress conditions Ssa4 is ∼60-fold less abundant than Ssa1 (Brownridge et al., 2013), points to functional specificity that is also consistent with specificity of genetic interactions, phenotypes, and transcript level changes in different *ssa* mutants (Hasin et al., 2014; Lotz et al., 2019).

As a way of explaining how slow-folding mutant septins are so efficiently excluded from assembly into septin complexes in cells also expressing a wild-type version of the same septin, we previously proposed that the G1 phase of the cell cycle represents a “window of opportunity” for septin complex assembly (Schaefer et al., 2016). According to our model, if a mutant septin has not already incorporated into septin hetero-octamers by the end of G1, it is unable to do so thereafter, and is found primarily in the cytosol/nucleus, presumably bound by chaperones (Johnson et al., 2015; Schaefer et al., 2016). Since we use cell divisions to mark time passed since synthesis, this model predicts that a slow-folding mutant septin would be unable to assemble post-translationally in our assay. Our results with Cdc3(Gly365Arg)-GFP were inconsistent with this prediction. We previously showed that the Cdc3 NTE protects another G-interface mutant, D289N, from being “outcompeted” by wild-type Cdc3 during assembly into septin complexes, likely because it shields the G interface from binding to specific chaperones that would otherwise inhibit interactions between the mutant Cdc3 and its G dimer partner, Cdc10 (Weems and McMurray, 2017). We note that while our split-YFP results pointed to a few differences in the way cytosolic chaperones interact with Cdc3 compared to the other septins, all bound to Ssb and at least one subunit of CCT (Denney et al., 2021). We interpret these differences in septin behavior as evidence that while special septin features like an NTE may alter interactions with some chaperones and influence the ability of a mutant septin to compete with its wild-type counterpart –– *i.e.*, in the context of “quality control” –– Ssb, CCT, and Ssa2-4 likely promote the folding of all wild-type septins.

Our *in vivo* crosslinking studies showing that chaperonins directly bind residues at or near the septin-septin G interface fit with our previous findings that mutations in or near the G interface generally prolong chaperone binding, presumably by delaying the acquisition of the native conformation capable of binding another septin and thereby rendering the sites inaccessible to chaperones. Since we did not test them, we cannot rule out other binding sites. It makes intuitive sense, however, that the septin NC interface, which at least for Cdc11 and Shs1 is exposed at both ends of a septin octamer, would not attract much cytosolic chaperone attention. A recent study that crosslinked Lys residues (Nitika et al., 2021) found a direct interaction between Ssa1 and a Lys (478) in the CTE of Cdc3, far from the G and NC interfaces (Supplemental Figure 9) and within the putative coiled coil formed in the non-native G Cdc3 homodimer (Figure 1). These crosslinks with Ssa1 probably only occurred because the other Ssa chaperones were eliminated (Nitika et al., 2021), otherwise Ssa2, Ssa3 or Ssa4 would presumably have “outcompeted” Ssa1 for septin interaction. As the crosslinked lysine in Ssa1 lies on the surface of the lid of the substrate-binding domain, it is possible that, once the Cdc3 NTE is displaced by interaction with Cdc12, the Ssa1 substrate-binding pocket does engage the Cdc3 G interface, bringing the Lys-rich Cdc3 CTE near the Ssa1 lid (Supplemental Figure 9).

In terms of CCT binding, if the client peptides bind in opposite orientations, the pattern of polar and hydrophobic residues in the Cdc10 sequence surrounding Trp255 (RGRKTRWSAINVED, in single-letter code), where Bpa substitution yielded crosslinks to Cct3, resembles the part of the HIV p6 protein (DKELYPLTSLRSLFGN) contact by the apical domain of human Cct3 (Joachimiak et al., 2014). Each CCT subunit has a distinct binding specificity (Balchin et al., 2018; Joachimiak et al., 2014) and, as indicated by our split-YFP studies (Denney et al., 2021), despite overall structural similarity different septins are likely contacted by different combinations of CCT subunits within the folding chamber. GroEL, too, relies on a distribution of polar and nonpolar surfaces to bind its clients (Chen and Sigler, 1999). Known sites of GroEL binding in other proteins (Wang et al., 1999) also match the highly hydrophobic sequence surrounding Cdc10(Tyr149Bpa) (Figure 2A), which also crosslinked to GroEL. Nonetheless, as with the Ssa1 example, it is possible that the GroEL crosslinks reflect “incidental” contacts away from the site(s) of direct chaperonin-Cdc10 interaction. More broadly, the overall three-layer α-β sandwich Rossmann fold of the septin GTPase domain matches the known preferences of both GroEL (Houry et al., 1999) and CCT (Stein et al., 2019). Indeed, an unbiased approach identified the GTPase EF-Tu, a known GroEL client *in vivo* (Houry et al., 1999), as the *E. coli* protein with highest similarity to all available septin structures (Supplemental Figure 10).

A homo-oligomeric Group II chaperonin can partially replace *E. coli* GroEL; substitution at a single residue even allows it to restore growth to *ΔgroEL*-mutant *E. coli* (Shah et al., 2016). Nonetheless, GroEL and CCT bind actin very differently (Balchin et al., 2018) and we want to be careful not to imply that the two chaperonins operate exactly the same way in septin folding. Septins may be among the proteins recently proposed to rely on CCT as a more GroEL-like “Anfinsen cage” that generally prevents aggregation and promotes native folding, without evolving a suite of specific contacts that establish client topology and direct a step-wise folding progression within the CCT chamber (Gestaut et al., 2022). GroEL also works differently than DnaK but each can support the folding of some clients in the absence of the other (Calloni et al., 2012). The ability of Δ*dnaK* and other *E. coli* chaperone mutants to make septin octamers does not rule out roles for those chaperones in septin folding; other chaperones may compensate for their absence. Supplementing the PURExpress reactions with additional purified chaperones might further increase Cdc12 yield. Ultimately, a septin’s path to the native conformation may look quite different depending on whether it passes through CCT or GroEL, even if the destination is the same. Finally, the ribosome itself acts as a chaperone to guide co-translational protein folding (Cassaignau et al., 2020), and we cannot exclude the possibility that differences between eukaryotic and prokaryotic ribosomes also create distinct folding trajectories with different chaperone requirements. Reconstituted eukaryotic cell-free translation systems to which purified chaperones can be added (Machida et al., 2021) may be especially useful in this regard.

While additional experiments will be required to identify exactly which features of Cdc12 confer its translational dependence on GroEL, we note that published polysome profiling experiments found that the density of ribosomes per 100 nucleotides on Cdc12 mRNA (0.87) is higher than on Cdc3 (0.45), Cdc10 (0.72), or Cdc11 (0.38) (Arava et al., 2003). Thus ribosomes move slowly along Cdc12 and may be on the verge of stalling. Single-molecule experiments show that the free energy drop accompanying co-translational folding can overcome ribosome stalling (Goldman et al., 2015). Closely-spaced ribosomes would put Cdc12 at highest risk for stall-induced collision and degradation of the mRNA and nascent protein when the pulling force from GroEL-assisted co-translational folding is lost. Like the “translational ramp” observed for many proteins in diverse species (Fredrick and Ibba, 2010), ribosome accumulation on the Pro-rich cluster at the Cdc12 N terminus likely acts to promote regular spacing of ribosomes downstream, preventing collisions. Eliminating the Pro-rich cluster probably tightens ribosome spacing on the main body of Cdc12 mRNA, favoring ribosome collisions when GroEL is even slightly dysfunctional. Indeed, a very recent study showed that a cluster of slowly-translated codons near the start codon stabilizes mRNAs with downstream slow-downs, presumably by preventing ribosome “traffic jams” (Sharma et al., 2021).

Others have noted that while “programmed” slow translation, such as by suboptimal codons, risks collision-induced mRNA and nascent protein degradation, it can also facilitate co-translational folding and assembly by providing more time for interactions to occur (Collart and Weiss, 2020). If high ribosome density on Cdc12 mRNA reflects especially slow translation even in wild-type cells, it is tempting to speculate that Cdc12 acts as a platform for co-translational septin assembly. Our previous studies (Weems and McMurray, 2017) support a model in which Cdc12 is the centerpiece for assembly of Cdc11/Shs1–Cdc12–Cdc3 trimers (Figure 6), which are subsequently brought together into hetero-octamers by a central Cdc10 homodimer. In this model, Cdc12 that has bound but not yet hydrolyzed GTP preferentially recruits Cdc11 (as opposed to Shs1), before interacting via its NC interface with Cdc3 (Weems and McMurray, 2017). Tubulins achieve a near-native conformation within the CCT chamber that is capable of binding GTP (Gestaut et al., 2022; Tian et al., 1995b). If it is translated slowly enough, quasi-native Cdc12•GTP might bind Cdc11 at its G interface and Cdc3 at its NC interface prior to ribosome release. The C-terminal ∼90 residues of Cdc12 are dispensable for stable hetero-octamer assembly (Bertin et al., 2010), so assembly could, in principle, be finished by the time Cdc12 translation is only ∼80% complete.

Like Cdc12 compared to other yeast septins, different human septins may differ in their translation speeds and the effects of variation in chaperone activity. Variation in chaperone expression between different cell types can also drastically alter the ability of a protein to fold, as illustrated by the example of an oncogenic p53 mutant that misfolds in other cell types but folds and functions normally in embryonic stem cells, where CCT and other chaperones are more highly expressed and bind the mutant protein (Rivlin et al., 2014). We previously showed how the presence of a “chemical chaperone”, guanidine, alters Cdc3 folding to promote a septin assembly pathway that bypasses incorporation of Cdc10 (Johnson et al., 2020). We think our new findings point to cytosolic chaperones as key factors in dictating the landscape of septin complexes assembled in a given cell type.

## MATERIALS AND METHODS

Detailed information about strains, plasmids, and oligonucleotides are provided in Supplementary Table 4.

### Microbial cultivation

Yeast were cultured using standard methods (Amberg et al., 2005) in liquid or solid (2% agar) rich (YPD; 1% yeast extract, 2% peptone, 2% glucose) or synthetic medium (SC; per liter, 20 g glucose, 1.7 g yeast nitrogen base without amino acids or ammonium sulfate, 5 g ammonium sulfate, 0.05 g tyrosine, 0.01 g arginine, 0.05 g aspartate, 0.05 g phenylalanine, 0.05 g proline, 0.05 g serine, 0.1 g threonine, 0.05 g valine, 0.05 g histidine, 0.1 g uracil, and 0.1 g leucine). Where appropriate to maintain plasmid selection, synthetic medium lacked specific components (uracil, tryptophan, histidine, etc.). Yeast were transformed using the Frozen-EZ Transformation II kit (Zymo Research # T2001). Bacteria were cultured in liquid or solid (1.5% agar) LB medium (1% tryptone, 1% NaCl, 0.5% yeast extract) containing appropriate antibiotics and additives needed for plasmid selection and induction of gene expression. Unless otherwise stated, antibiotics and arabinose were used at the following final concentrations from 1000x stocks dissolved in water: carbenicillin at 70 µg/mL, ampicillin at 100 µg/mL, kanamycin at 40-50 µg/mL, chloramphenicol at 34 µg/mL (stock dissolved in ethanol), spectinomycin at 50 µg/mL, arabinose at 0.02%. 1 M p-benzoyl-l-phenylalanine (Bpa, Advanced Chemtech #YF2424) solution was freshly made before use by dissolving 27 mg Bpa in 100 µL 1 M NaOH. Bacterial transformation was performed using the Mix & Go! *E. coli* Transformation Kit (Zymo Research #T3001) or by making cells chemically competent using CaCl_2_ (Sambrook and Russell, 2006).

### Lysogenization and construction of *E. coli* chaperone deletion mutants

Lysogenization was accomplished with the λDE3 Lysogenization Kit (Novagen #69734- 3). 1-2 µL host cells grown to mid-log phase at 37°C in LB containing 0.2% maltose and 10 mM MgCl_2_ were combined with the three λ phages (10^8^ CFU each), incubated at 37°C for 20 min, and spotted to LB. Several dozen to several hundred λDE3 lysogens were obtained following overnight growth at 37°C for strains BW25113, JW2573, and JW0462, which are from the “Keio” deletion collection (Baba et al., 2006). Candidate lysogens were assayed with tester phage in top agarose (1% tryptone, 0.5% NaCl, 0.6% agarose), where plaques formed in the presence of 0.4 mM IPTG were slightly larger than those without IPTG. Lysogens were further confirmed by their immunity to selection phage and by visualization of a T7-driven GFP reporter (encoded by plasmid pDual-eGFP; Supplemental Figure 11). Recombineering was accomplished using arabinose-induced expression of λ-red recombination genes from pSIJ8 (Jensen et al., 2015) maintained in host cells prior to transformation with a *dnaJ*- or *dnaK*-targeting kanamycin resistance amplicon from strain JW0014 using dnaJ_keio_fw and dnaJ_keio-re, or from strain JW0013 using dnaK_keio_fw and dnaK_keio_re. Bacteria were made chemically competent for recombineering using the Z-competent™ kit (Zymo Research # T3001) and recovered in SOC at 30°C overnight before plating on solid LB containing kanamycin at 30°C. The presence of the kanamycin resistance cassette insertion and the absence of duplicated wild-type allele were verified by PCR (Supplementary Figure 11). The temperature-sensitive loss of pSIJ8 was subsequently confirmed in recombinants. Note that host cells lacking DnaJ and DnaK cannot be lysogenized with λDE3 owing to the essential functions of those gene products in lambda phage replication. For this reason, Δ*dnaJ* and Δ*dnaK* were recombineered *de novo* in lysogen strain B002.

### *In vivo* photocrosslinking

Yeast cells of the *upf1Δ* strain YRP2838 carrying pSNRtRNA-pBPA-RS (Krishnamurthy et al., 2011) and a plasmid encoding a GST-6xHis-tagged septin were inoculated from single colonies into 5 mL liquid synthetic medium lacking tryptophan and uracil and containing 2% raffinose. Septin expression plasmids were from a collection encoding GST-6xHis-tagged proteins (Zhu et al., 2001), or mutant derivatives thereof. Following overnight growth at 30°C, the cultures were pelleted at 3,500 x *g* for 5 min, washed once in fresh medium, and resuspended in a 250-mL baffled shaker flask at OD_600_=0.03 in 50 mL of fresh liquid synthetic medium lacking tryptophan and uracil and containing 2% raffinose. Cultures were grown shaking at 225 rpm and 30°C to OD_600_=0.5, at which point 50 µL of a 1 M solution of Bpa was added dropwise to the flask while swirling. Galactose was immediately added to 2% final concentration, and the cultures were rotated at 30°C for another 3 hours. Cells were exposed, or not, to 254-nm light for 30 minutes in a Chromato-vue cabinet (Model C-70, Ultra-Violet Products, Inc.) for 30 minutes in flat-bottom 6-well dishes. Cultures were centrifuged at 4,000 rpm, washed once with Equilibration Buffer A (300 mM NaCl, 50 mM Na_2_PO_4_, 50 mM Tris-HCl pH 8.0, 0.05% octylphenoxy poly(ethyleneoxy)ethanol [Nonidet P-40 equivalent; Millipore-Sigma #i8896], 8 M urea) containing protease-inhibitor mix (Complete EDTA-free; Roche # 11 873 580 001), and frozen at -70°C. For detection of crosslinked species, cell pellets were resuspended in 400 µL Equilibration Buffer A containing protease-inhibitor mix (Complete EDTA-free; 11 873 580 001, Roche). Cells were lysed using the equivalent of 100 µL glass beads and 4 cycles of 30-second on/off vortexing (incubating on ice during the “off” cycle). Samples were centrifuged at maximum speed to remove glass beads. Another 400 µL Equilibration Buffer A was added to bead-free lysate and lysates were clarified by centrifugation at 12,000 x g at 4°C for 15 minutes. Clarified lysates were added to previously equilibrated His-Pur™ Ni-NTA spin columns (0.2 mL resin bed, Thermo Scientific #88224) and incubated at room temperature for 2 hours. Flow-through (“Unbound”) was transferred to another tube and columns washed twice, first with Equilibration Buffer A and second with Wash Buffer C (300 mM NaCl, 50 mM Na_2_PO_4_, 50 mM Tris-HCl pH 8.0, 0.05% octylphenoxy poly(ethyleneoxy)ethanol, 8 M urea, 10 mM imidazole). His-tagged proteins (“Bound”) were eluted in 200 µL Elution Buffer B (300 mM NaCl, 50 mM Na2PO4, 50 mM Tris-HCl pH 8.0, 0.05% octylphenoxy poly(ethyleneoxy)ethanol, 8 M urea, 250 mM imidazole).

For crosslinking in *E. coli*, single colonies of strain BL21(DE3) carrying plasmid pEVOL-pBpF (Chin et al., 2002) and a pET21-based Cdc10-encoding plasmid were used to inoculate 5-mL liquid cultures of LB with ampicillin and chloramphenicol. Following overnight growth at 37°C, the cultures were pelleted at 3,500 x *g* for 5 minutes at room temperature, washed once in fresh LB with antibiotics, and resuspended in a 250-mL baffled shaker flask at an OD_600_=0.03 in 50 mL of fresh LB with antibiotics plus 0.02% arabinose to induce expression of the modified tRNA. Cultures were shaken at 37°C until the OD_600_ reached 0.5, at which point 50 µL of a 1 M solution of Bpa (made fresh by dissolving 27 mg in 100 µL 1 M NaOH) was added dropwise to the flask while swirling. IPTG was immediately added to 0.3 mM final concentration, and the cultures were shaken at 37°C for another 3 hours. Cells were exposed, or not, to 254-nm light for 30 minutes as described above. Cultures were centrifuged at 4000 rpm, washed once with Equilibration Buffer B (300 mM NaCl, 20 mM Na_2_PO_4_, 8 M urea, 10 mM imidazole; pH 7.4) containing protease-inhibitor mix (Complete EDTA-free; 11 873 580 001, Roche), and frozen at -70°C. For detection of crosslinked species, cell pellets were resuspended in 1 mL Equilibration Buffer B containing protease-inhibitor mix (Complete EDTA-free; 11 873 580 001, Roche) and 0.2 mg/mL lysozyme. Cells were incubated on ice for 15 min and subjected to two 15-sec pulses of sonication (550 Sonic Dismembrator, Thermo Scientific) to complete lysis. Lysates were clarified by centrifugation at 12,000 x g at 4°C for 15 min. Clarified lysates were added to previously equilibrated His-Pur™ Ni-NTA spin columns (0.2 mL resin bed, Thermo Scientific #88224) and mixed end-over-end at 4°C for 30 min. Flow-through was discarded and columns washed twice with Wash Buffer D (300 mM NaCl, 20 mM Na_2_PO_4_, 8 M urea, 25 mM imidazole; pH 7.4). His-tagged proteins (“Bound”) were eluted in 200 µL Elution Buffer C (300 mM NaCl, 50 mM Na_2_PO_4_, 50 mM Tris-HCl pH 8.0, 0.05% octylphenoxy poly(ethyleneoxy)ethanol, 8 M urea, 250 mM imidazole). Bound fractions were combined 1:1 with 2x SDS–PAGE sample buffer containing 10 mM DTT and heated at 95°C for 5 min prior to SDS-PAGE.

### Heterologous co-expression of yeast septins

Cells carrying relevant plasmids described in figure legends were inoculated from single colonies into 3-mL LB cultures with indicated antibiotics and 0.02% arabinose (none for “No Septin Control”, ampicillin and chloramphenicol for B002, B003, B004, B005, and B006; ampicillin, chloramphenicol, and spectinomycin for strain 461). Cultures were grown overnight at 24°C then pelleted and resuspended in fresh LB, after which 1-2 mL was inoculated in a 250-mL baffled shaker flask containing 200 mL fresh LB with indicated antibiotics and 0.02% arabinose. Cultures were shaken at 225 rpm and 24°C to OD_600_=0.6, after which 100 mL was removed and placed in a new flask and grown shaken at 37°C. Both 100-mL cultures were induced with 0.3 mM final IPTG, the original culture immediately at 24°C and the new culture following a 15-min acclimation period at 37°C. After 3 hours cultures were centrifuged at 4000 rpm, washed once with Lysis Buffer B (300 mM NaCl, 2 mM MgCl_2_, 1 mM EDTA, 5 mM β-mercaptoethanol, 0.5% Tween 20, 12% glycerol, 50 mM Tris-HCl, pH 8), and frozen at -70°C.

### Expression and purification of individual yeast septins

6xHis-Cdc3 and Cdc12 were individually expressed in *E. coli* and purified by Keyclone Technologies. 6xHis-Cdc3 was produced using plasmid pBEG2 (Versele and Thorner, 2004). For Cdc12, a plasmid was constructed (pMBP-His-Cdc12) that fused to the N terminus of Cdc12 maltose-binding protein (MBP), a heptahistidine (7xHis) tag, and a tobacco-etch mosaic virus (TEV) protease cleavage site. 300 mL of BL21(DE3) cells carrying the appropriate plasmid was induced at OD_600_ of 0.6 with 0.3 mM IPTG for 4 hr at 32°C with shaking at 120 rpm. Cells were pelleted and lysed by sonication. The lysate was clarified by centrifugation and the supernatant was loaded on a Ni-NTA column (ThermoFisher Scientific) followed by washing and elution. The MBP-7xHis portion was cleaved using 6xHis-tagged TEV enzyme made by Keyclone Technologies, and the MBP-7xHis and 6xHis-TEV enzyme were removed by passing through the Ni-NTA column again after enzyme cleavage.

### Mass spectrometry to identify yeast or *E. coli* proteins in crosslinking and *in vitro* translation experiments

Analysis was performed by the Mass Spectrometry Proteomics Shared Resource Facility of the University of Colorado Anschutz Medical Campus. Samples were processed with a filter-aided sample preparation (FASP) protocol (Wiśniewski, 2016). Briefly, aliquots containing 36 μg of total protein were mixed with 200 μL of 8M urea in 0.1M (ABC) ammonium bicarbonate (pH 8.5) reduced with 5 mM TCEP (tris(2-carboxyethyl)phosphine) for 20 min and alkylated with 50 mM 2-chloroacetamide for 15 min in the dark, all at room temperature. Reduced and alkylated samples were loaded onto 10 kDa cut-off filters (Vivacon 500, Sartorius #VN01H02) and centrifuged at 14,000 × g for 20 min. The samples were then washed twice with 50 mM ABC and trypsin was added at an enzyme:protein ratio of 1:20 for overnight digestion at 37°C. Following trypsin digestion, samples were centrifuged at 14,000 × g for 20 min at room temperate and acidified with formic acid (FA) to a final concentration of 0.1%. Aliquots containing 10 μg of digested peptides were purified using PierceTM C18 Spin Tips (Thermo Scientific # 84850) according to the manufacturer’s protocol, dried in a vacuum centrifuge, and resuspended in 0.1% FA in mass spectrometry-grade water. For crosslinking experiments, digested peptides were subjected to liquid chromatography mass spectrometry (LC-MS/MS) using an Eksigent nanoLC Ultra coupled to a LTQ-Orbitrap Velos Pro mass spectrometer. Fragmentation spectra were interpreted against the UniProt Fungi database using the Mascot search engine. One missed tryptic cleavage was allowed, and the precursor-ion mass tolerance and fragment-ion mass tolerance were set to 15 ppm and .6 Da, respectively. Carbamidomethyl (C) was selected as a fixed modification and oxidation (M) was selected as a variable modification. The protein-level FDR was ≤ 1%. For in vitro translation experiments, LC-MS/MS was performed using an Easy nLC 1000 instrument coupled to a Q Exactive™ HF Mass Spectrometer (both from ThermoFisher Scientific). Peptides were loaded on a C18 column (100 μM inner diameter x 20 cm) packed in-house with 2.7 μm Cortecs C18 resin, and separated at a flow rate of 0.4 μl/min with solution A (0.1% FA) and solution B (0.1% FA in acetonitrile) and under the following conditions: isocratic at 4% B for 3 minutes, followed by 4%-32% B for 102 minutes, 32%-55% B for 5 minutes, 55%-95% B for 1 min and isocratic at 95% B for 9 minutes. The mass spectrometer was operated in data-dependent acquisition (DDA) mode with the top 15 most abundant precursors being selected for MS/MS analysis. Fragmentation spectra were searched against the UniProt Escherichia coli proteome database combined with the Cdc10 and Cdc12 protein sequences using the MSFragger-based FragPipe computational platform (Kong et al., 2017). Contaminants and reverse decoys and were added to the database automatically. The precursor-ion mass tolerance and fragment-ion mass tolerance were set to 10 ppm and .2 Da, respectively. Fixed modifications were set as carbamidomethyl (B) and oxidation (M) was selected as a variable modification. Two missed tryptic cleavages were allowed, and the protein-level false discovery rate (FDR) was ≤ 1%.

### Hydrogen-deuterium exchange mass spectrometry

HDX-MS was performed at the University of Colorado, Boulder Central Analytical Mass Spectrometry lab. To identify peptic fragments of Cdc3, 100 pmoles of protein (5 μL) were mixed with 45 μL of the sample buffer (150 mM NaCl, 25 mM HEPES, 5% glycerol, pH 7.4) and then mixed with 50 μL of a quenching buffer (3 M guanidine HCl, 1.5% formic acid). The whole sample (100 μL total) was injected into a Waters HDX-LC box (Waters) in which protein was digested by an online pepsin column (Waters Enzymate BEH pepsin column, 2.1 × 30 mm, 5 μm and the resulting peptic peptides were trapped and desalted on a Waters ACQUITY UPLC BEH C4 1.7 μm VanGuard Pre-column (2.1 × 5 mm) at 100 µL/min Buffer A (0.1% formic acid in water) for 3 min. The digestion chamber was kept at 15 °C and the trap and analytical columns were at 0°C. The peptides were eluted with 3-33 % Buffer B (0.1% formic acid in acetonitrile) between 0-6 min, 33-40% B between 6-6.5 min, and 40-85% B between 6.5-7 min. The eluted peptides were resolved a Waters ACQUITY UPLC Protein BEH C4 column (300 Å, 1.7 μm, 1 mm × 50 mm). MS/MS spectra were performed on a Thermo LTQ orbitrap Velos mass spectrometer. The peptides were ionized using electrospray ionization (ESI) with the source voltage = 5.0 kV and S-lens RF level = 60%. The capillary temperature was 275 °C and the source heater was off. The sheath gas flow was 10. Precursor ions were scanned between 350-1,800 m/z at 60,000 resolution with AGC 1 × 10^6^ (max ion fill time = 500 ms). From the precursor scan, the top 10 most intense ions were selected for MS/MS with 180 s dynamic exclusion (10 ppm exclusion window, repeat count = 1) and AGC 1 × 10^4^ (max ion fill time = 100 ms) Ions with unassigned charge states were rejected for MS/MS. The normalized collision energy was 35%, with activation Q = 0.25 for 10 msec. MS/MS spectra were searched against a database consisting of Cdc3 protein sequence in a FASTA format using the MaxQuant/Andromeda program (version 1.6.3.4) developed by the Cox Lab at the Max Planck Institute of Biochemistry (Cox and Mann, 2008; Cox et al., 2011). The digestion mode was “unspecific” and oxidation of methionine was set as a variable modification. The minimum peptide length for the unspecific search was 8. MaxQuant/Andromeda used 4.5 ppm for the main search peptide tolerance and 0.5 Da for MS/MS tolerance. The false discovery rate was 0.01. A list of peptides identified from the search was used as an exclusion list for the next round of LC-MS/MS and a total of three LC-MS/MS were performed for each protein.

For hydrogen/deuterium exchange, ten 250 µL aliquots of the sample buffer were dried using vacuum centrifugation, reconstituted with an equal volume of deuterium oxide (“D_2_O buffer”), and put in a 10°C water bath. Ten minutes prior to the initiation of HDX reaction, 5 µL of Cdc3 protein (100 pmoles) was transferred to a 0.5 mL tube and incubated at 10 °C. HDX reaction was initiated by the addition of the D_2_O buffer (45 µL) to the 10 µL sample (90% D_2_O final). The reaction was quenched at 15 sec, 30 sec, 1 min, 2 min, 5 min, 10 min, 20 min, 40 min, 80 min, and 120 min after the initiation by the addition of the quench buffer (50 µL). The whole sample (100 µL) was injected into the Waters HDX-LC box and LC-MS was performed as described above. For HDX samples, only MS1 spectra were recorded. For the 0 sec control, 100 pmoles of Cdc3 (5 µl) was mixed with 50 µL of the quenching solution, and then with 45 µL of the D_2_O buffer. HDX-MS data were analyzed using Mass Spec Studio (version 2.4.0.3484) developed by the Schriemer lab at the University of Calgary (Raval et al., 2021; Rey et al., 2014). “Peptide.txt” and “evidence.txt” from the MaxQuant/Andromeda search results were used to generate the “identification” table for Mass Spec Studio using an in-house Python script. The raw files from the Thermo LTQ orbitrap Velos were converted to mzML files using ProteoWizard (version 3.0.20216, 64 bit). The default processing parameters were used except mass tolerance = 15 ppm, total retention time width = 0.15 min, XIC smoothing = Savitzky Golay Smoothing, and deconvolution method = centroid. All processed data were manually validated. Exchange ratio values were mapped onto the AlphaFold2 predicted Cdc3 structure using the “spectrum b” function in the PyMOL Molecular Graphics System v2.0 (Schrödinger, LLC). Briefly, the .pdb file was edited to replace the B values with average exchange ratios at 120 minutes. Residues not detected by mass spectrometry were assigned an exchange value of 2 and the structure rendered in PyMol was colored using the command “spectrum b, blue red grey, minimum=0, maximum=2”.

### ATPase and refolding activity of purified GroEL/ES

GroES and GroEL were obtained from Takara Biotechnology (Dalian) Co., Ltd (#7331 and #7330). ATPase activity was measured using the EnzChek™ Phosphate Assay Kit (ThermoFisher Scientific # E6646) according to the manufacturer’s instructions with 500 nM GroEL/ES tetradecamer. Rhodanese refolding was monitored as absorbance at 460 nm in a 96-well plate using the same device. Rhodanese (Sigma #R1756-5MG) was denatured in 8 M urea. The thiocyanate enzymatic assay stock was 140 mM KH_2_PO_4_, 160 mM KCN (Sigma-Aldrich #207810), and 160 mM Na_2_S_2_O_3_ in water. Reagent concentrations during refolding reactions were: 300 nM GroEL, 360 nM GroES, 250 nM denatured Rhodanese, 1 mM ATP. Formaldehyde quench stock was 30% formaldehyde in water. Ferric nitrate reporter stock was 8.5% w/v Fe(NO_3_)_3_ and 11.3% v/v HNO_3_ in water. Absorbance at 360 nm was measured after 15 minutes using a 96-well plate using a plate reader (Cytation 3, BioTek).

### *In vitro* translation

The PURExpress ΔRF123 kit (New England Biolabs #E6850S) was used with the following modifications from the manufacturer’s instructions. As described previously (Kawahashi et al., 2007), fluorescent puromycin (6-FAM-dC-Puromycin, Jena Bioscience #NU-925-6FM-L) was added to 4.6 µM. Purified EF-P was purchased from Biomatik (#RPC22718) and, as described in (Ude et al., 2013), was added to 1 µM where indicated, or a control buffer (20 mM Tris-HCl pH 8.0, 0.5 M NaCl, 6% Trehalose) was added. Purified GroES and GroEL were each added to 0.87 µM where indicated, or a control buffer (50 mM Tris-HCl pH 7.4, 50 mM KCl, 10 mM MgCl_2_) was added. Additional ATP was added to 6.7 mM (not including ATP included in PURExpress kit reagents) to prevent ATP depletion resulting from added GroEL/ES. 62 ng of template plasmid (pMVB128) was added to each reaction, and 16 U of RNase inhibitor was included. Per manufacturer’s instructions, the reactions were completed at 37°C, and the total reaction time was 3 hours. Completed reactions were analyzed immediately or stored at –20C.

### Analysis of purified septin oligomerization state

Purified Adh1 (Sigma-Aldrich #A8656-1VL) or Cdc3 were diluted in PBS and centrifuged through 0.5 mL 100-kDa molecular-weight cutoff protein concentrators (Thermo Scientific Pierce #PI88503) according to the manufacturer’s instructions. Analytical SEC was performed with an Agilent AdvanceBio SEC 300Å 2.7 µm w/Guard Column equilibrated and run at 0.3 mL/min in PBS with GTP (Sigma-Aldrich #51120) or guanidine added as indicated. Protein was injected in 50 µL. Molecular weight standards were AdvanceBio SEC 300A Protein Standard (#5190-9417).

### SDS–PAGE and immunoblotting

Where indicated in figure legends, proteins were extracted from yeast cells using an alkaline lysis/trichloroacetic acid (TCA) precipitation method (Hase et al., 1983). Proteins were typically resolved on 10% SDS-PAGE gels. Immunoblotting was performed via semi-dry transfer (Bio-Rad #1703940) to PVDF membranes. Primary antibodies: anti-GroEL (Enzo Life Sciences #ADI-SPS-870-D, RRID:AB_2039163), anti-DnaK (Enzo Life Sciences #ADI-SPA-880-F, RRID:AB_10619012), anti-GST (Santa Cruz Biotechnology #sc-459, RRID:AB_631586), anti-6xHis (UBPBio # Y1011), anti-Cdc11 (Santa Cruz Biotechnology # sc-7170, RRID:AB_671797), anti-Cdc3 (McMurray and Thorner, 2008). Secondary antibodies: Infrared-labeled anti-mouse (680 nm, Biotium # 20061-1, RRID:AB_10854088), infrared-labeled anti-rabbit (800 nm, Cell Signaling Technology # 5151, RRID:AB_10697505). Immunoblots were scanned on a Li-Cor Odyssey. Gels with fluorescent puromycin were scanned on a Sapphire Biomolecular Imager (Azure Biosystems).

### *In vivo* assays of kinetics of septin folding and post-translational assembly

All experiments were carried out at room temperature (∼22°C). With a few exceptions, noted below and in the figure legends, cells were BY4741 (haploid wild-type parent strain) or BY4743 (diploid wild-type parent strain) or derivatives thereof carrying the plasmid(s) pTS395 (*URA3 P_GAL_–GFP*) (Carminati and Stearns, 1997), pMVB1 (*URA3 P_GAL_–CDC3-GFP*) (Schaefer et al., 2016), pMVB2 (*URA3 P_GAL_–CDC12-GFP*) (Versele et al., 2004), pMR287-D36N (*TRP1 P_GPD1_–YDJ1(D36N)*) (Reidy et al., 2014), YCpL-SSA2(R169H) (*LEU2 SSA2(R169H)*), pRS315 (*LEU2*) (Sikorski and Hieter, 1989), pRS314 (*TRP1*) (Sikorski and Hieter, 1989), YEpPgal–GST-His6-Cdc12 (*URA3 P_GAL_– GST-5xHis-CDC12*) (Zhu et al., 2001), and/or pGF-IVL-470 (*URA3 P_GAL_–CDC12-mCherry*). The *cct4* and *cdc3(G365R)* strains were from a BY4741-based temperature-sensitive mutant collection (Li et al., 2011). The *hsc82Δ* and *hsp82Δ* strains were from the deletion collection made in BY4741 (Winzeler et al., 1999). For experiments with *TRP1*-marked plasmids, the *trp1Δ* derivative of BY4741 was used. The *ssa1Δ* and *ssa4Δ* strains were from the deletion collection made in BY4742 (*MAT⍺ his3Δ leu2Δ lys2Δ ura3Δ*) (Winzeler et al., 1999). Cells containing plasmids were cultured to mid-log phase in synthetic complete media lacking uracil, leucine, and/or tryptophan and containing 2% raffinose or sodium lactate, which neither induces nor represses the *GAL1/10* promoter. The kinetics-of-folding experiment in *cct4* cells and the corresponding wild-type control experiment was performed by loading cells into an ONIX EV-262 Microfluidic Perfusion Platform (CellAsic #NX-262, plate #Y04C). Galactose-containing medium was flowed into the chamber and flow was maintained at a constant rate thereafter. To avoid phototoxicity, different cells were visualized at each timepoint after galactose addition, with the exception of several fields of cells that were imaged by transmitted light both at time 0 and after 13.5 hr for the purpose of retrospectively assessing relative division rate. For other kinetics-of-folding experiments, galactose was added at “time 0” to 0.05% or 0.1% final concentration to liquid raffinose or lactate cultures, and aliquots were removed at each timepoint and spotted onto agarose pads for microscopy. For post-translational assembly experiments, expression was terminated after 6 hours of induction by transferring cells to fresh medium containing 2% glucose. Aliquots of cells from glucose cultures were imaged at various timepoints and the number of cell divisions since the previous timepoint was calculated using a hemacytometer. Cultures were occasionally back-diluted into fresh media to prevent culture saturation, and cultures were temporarily stored overnight at 4°C as necessary throughout the data collection period.

### RT-PCR

500 µL of culture were preserved and pelleted with RNAprotect Bacteria Reagent (Qiagen #76506) before freezing at -70°C. Cell pellets were thawed and lysed in 200 µL TE (10 mM Tris-HCl, 1 mM EDTA, pH 8) buffer containing 20 mg total Proteinase K and 15 mg/mL lysozyme. RNA was isolated from bacterial lysate using RNeasy® Mini kit (Qiagen #74104) and eluted in sterile RNase-free water. RNA samples were quantified by Nanodrop and treated with TURBO DNase (Thermo Scientific Invitrogen #AM1907). ≤500 ng DNase-treated RNA was used to generate cDNA using Protoscript II Reverse Transcriptase (New England Biolabs #M0368) and the standard first-strand cDNA synthesis protocol. RT-PCR with OneTaq (New England Biolabs #M0480S) was performed on cDNA preparations, typically using 0.25-0.5 µL cDNA input in a 25-µL reaction with 25 cycles. Annealing temperature and extension time was 53°C and 50 seconds for Cdc12 and 51°C and 60 seconds for Cdc3.

### Microscopy

Images were captured with an EVOSfl all-in-one epifluorescence microscope (Thermo-Fisher Scientific) with a 60x oil objective and GFP (#AMEP4651) and Texas Red (#AMEP4655) LED/filter cubes. Image adjustment and analysis was carried out using FIJI software (Schindelin et al., 2012). Bud neck fluorescence was assessed with line scans (width 8 pixels) perpendicularly across bud necks measuring the difference between maximum and minimum pixel intensity values. The integrated pixel intensity inside an 8- or 10-pixel diameter circle was used to approximate cytoplasmic/nuclear fluorescent signal for each cell (vacuoles were avoided when possible). A minimum of 30 cells were analyzed at each timepoint.

### *In vivo* imaging of septin foci resulting from overexpression

JTY3992 (McMurray et al., 2011b) is derived from BY4742 with mCherry fused to the C terminus of Cdc10. The *HSP104-mCherry* strain was made by homologous recombination using a donor PCR product made with template plasmid pDK305 (Kaganovich et al., 2008) and primers GFPtomCherryfw and GFPtomCherryre70, co-transformed with plasmid pGF-V2226 (Roggenkamp et al., 2017) into cells of the *HSP104-GFP* strain from the BY4741-based GFP collection (Huh et al., 2003) that were already carrying plasmid pML104 (Laughery et al., 2015). Cas9-mediated cutting of the GFP sequence was repaired by homologous recombination with the mCherry-encoding PCR product, replacing the GFP and the downstream *HIS3MX* marker. The Cdc3-mCherry strain was made by integrating *Bgl*II-digested plasmid yIp128-Cdc3-mcherry (Tong et al., 2007) into BY4743. Chaperone deletion strains were from the BY4742- based haploid or BY4743-based homozygous diploid deletion collections (Winzeler et al., 1999). Cdc12-GFP foci were induced by culturing cells carrying pMVB2 in synthetic medium with 2% galactose at room temperature. Cdc12-mCherry was overexpressed in the same way using plasmid pGF-IVL-470. Cdc10(D182N)-GFP foci were induced by culturing cells carrying pPmet-cdc10-1-GFP in synthetic medium lacking methionine (to induce the *MET15* promoter) at room temperature or shifted to 37°C for 6 hours. For analysis of Cdc12-GFP focus appearance and disappearance in *hsp104Δ* mutants, synthetic cultures grown in 2% raffinose overnight were diluted into 25 mL fresh medium with 0.1% galactose and 1.9% raffinose. After 6 hours of induction, the cells were pelleted and resuspended in synthetic medium with 2% glucose.

### Predicted protein structures

AlphaFold predicted protein structures were accessed from the AlphaFold Protein Structure Database (Jumper et al., 2021; Varadi et al., 2022) at the following entries: Cdc3 (P32457), Cdc11 (P32458), Cdc12 (P32468), Ssa1 (P10591), Ssa2 (P10592), Ydj1 (P25491), and Zuo1 (P32527). Structures were imaged in PyMOL (Schrödinger LLC).

### Structural similarity search

The VAST+ server (Vector Alignment Search Tool +, https://www.ncbi.nlm.nih.gov/Structure/vastplus/vastplus.cgi) (Madej et al., 2014) was used to search for *E. coli* proteins with highest similarity to all septin structures available in the Protein Data Bank as of July 2020.

### Analysis of Cdc3 and Cdc12 sequence for potential translation-slowing properties

Data for codon adaptation index (CAI) for the Cdc3 and Cdc12 coding sequences were obtained from the Codon Usage Database at (https://www.kazusa.or.jp/codon/) for yeast (species 4932) and *E. coli* (species 83333) (Nakamura et al., 2000). For predicted mRNA secondary structures, the coding sequences were analyzed at the mFold RNA server (http://www.unafold.org/mfold/applications/rna-folding-form.php) (Zuker, 2003) using default settings/parameters. For amino acid pairs, each pair of amino acids in the Cdc3 and Cdc12 sequences was assessed according to published semi-quantitative bins of effect on translation elongation speed (Ahmed et al., 2020) and color-coded using a custom macro in Microsoft Excel.

## ACKNOWLEDGEMENTS

We thank Corrine Tuckey of New England Biolabs for insights and advice with PURExpress reactions, Thomas Lee of the Boulder Central Analytical Mass Spectrometry lab for carrying out the HDX-MS analysis, Monika Dzieciatkowska and Anthony Saviola of the Proteomics Mass Spec Facility for their MS analysis, Jun Yang of Keyclone Technologies for mutagenesis, cloning and protein expression and purification, Kevin Stein of the Frydman lab at Stanford for sharing unpublished results about co-translational interactions between chaperones and septins, and Greg Finnigan of Kansas State University for sharing unpublished septin and CRISPR plasmids. The work was supported by the National Institute of General Medical Sciences of the National Institutes of Health under Award R01GM124024 (to M.A.M.) and Award T32GM008730 (to A.S.D).

---

**Supplemental Figure 1.**
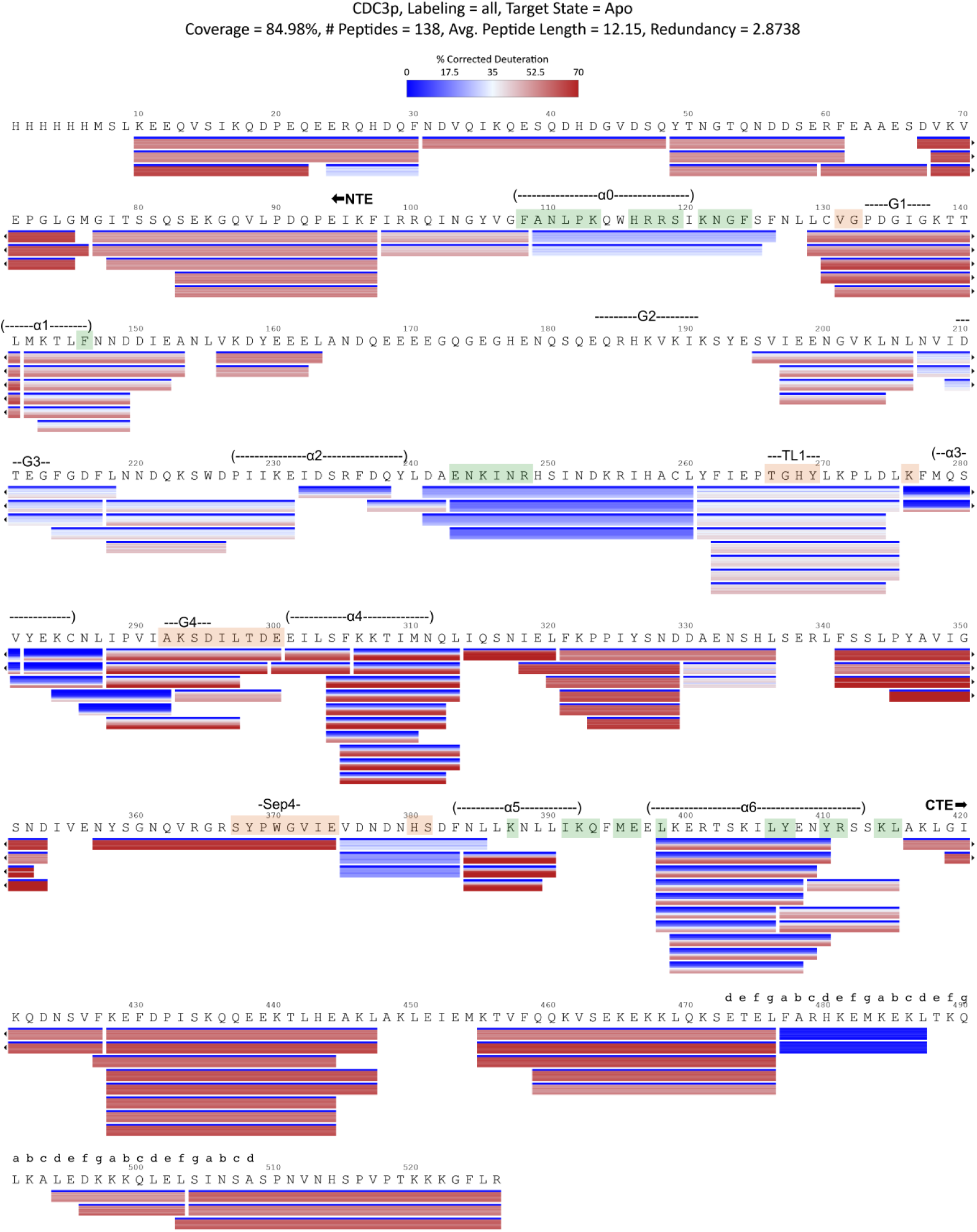
Hydrogen-deuterium exchange ratios for purified 6xHis-Cdc3 mapped onto its primary sequence. Mass Spec Studio (Rey et al., 2014) was used to display sequence coverage and deuteration values for 6xHis-Cdc3 sampled after 15, 30, 60, 120, 300, 600, 1200, 2400, 4800, and 7200 seconds in deuterated buffer. As indicated in the heat map key at top, horizontal lines underneath the protein sequence show peptide deuteration values with timepoints stacked vertically on each other. Orange or green highlighting and text above each line of sequence indicate points of contact across the G or NC homodimer interface, respectively, for the human septin SEPT2 (Sirajuddin et al., 2007). “abcdefg” indicate individual amino acids with predicted properties that follow the order expected for “hydrophobic heptad repeats”, as predicted by the PARCOILS2 algorithm (McDonnell et al., 2006). All annotated residues have PARCOILS2 probability >0.98 of forming a homotypic coiled coil.

**Supplemental Figure 2.**
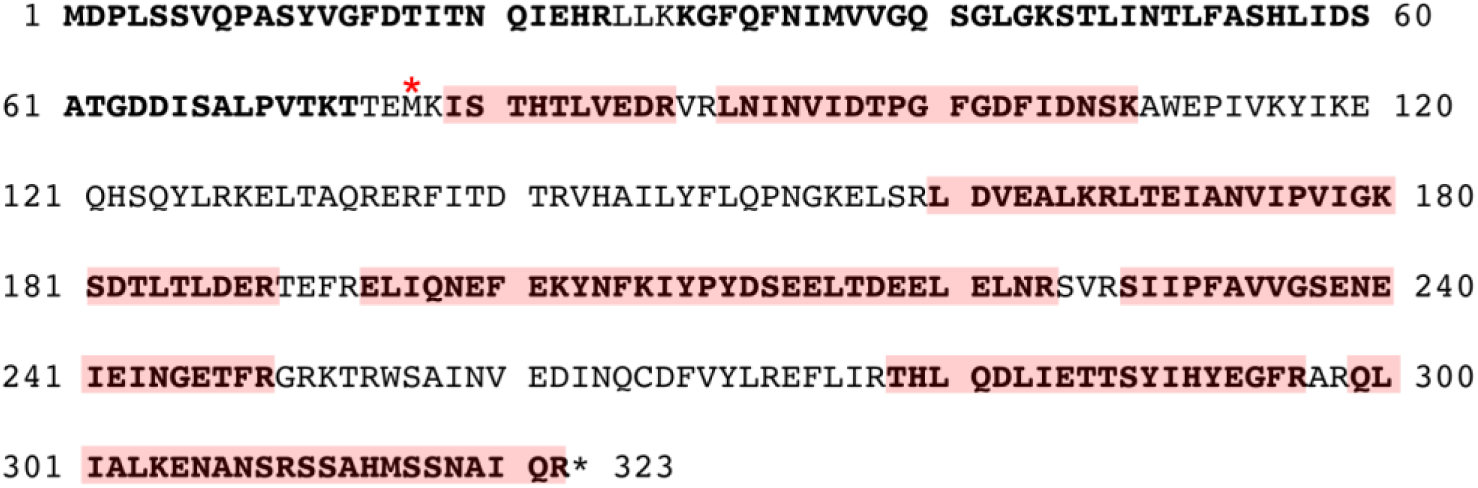
Mass spectrometry coverage of Cdc10 from excised bands in Figure 2D. Cdc10 protein sequence is shown with peptides detected by mass spectrometry. Bold, from bands migrating in SDS-PAGE above ∼37 kDa in Figure 2D, as expected for full-length Cdc10-6xHis; highlighted in red, band migrating at ∼25 kDa. Red asterisk is over the Met residue corresponding to the presumptive start codon for the truncated protein.

**Supplemental Figure 3.**
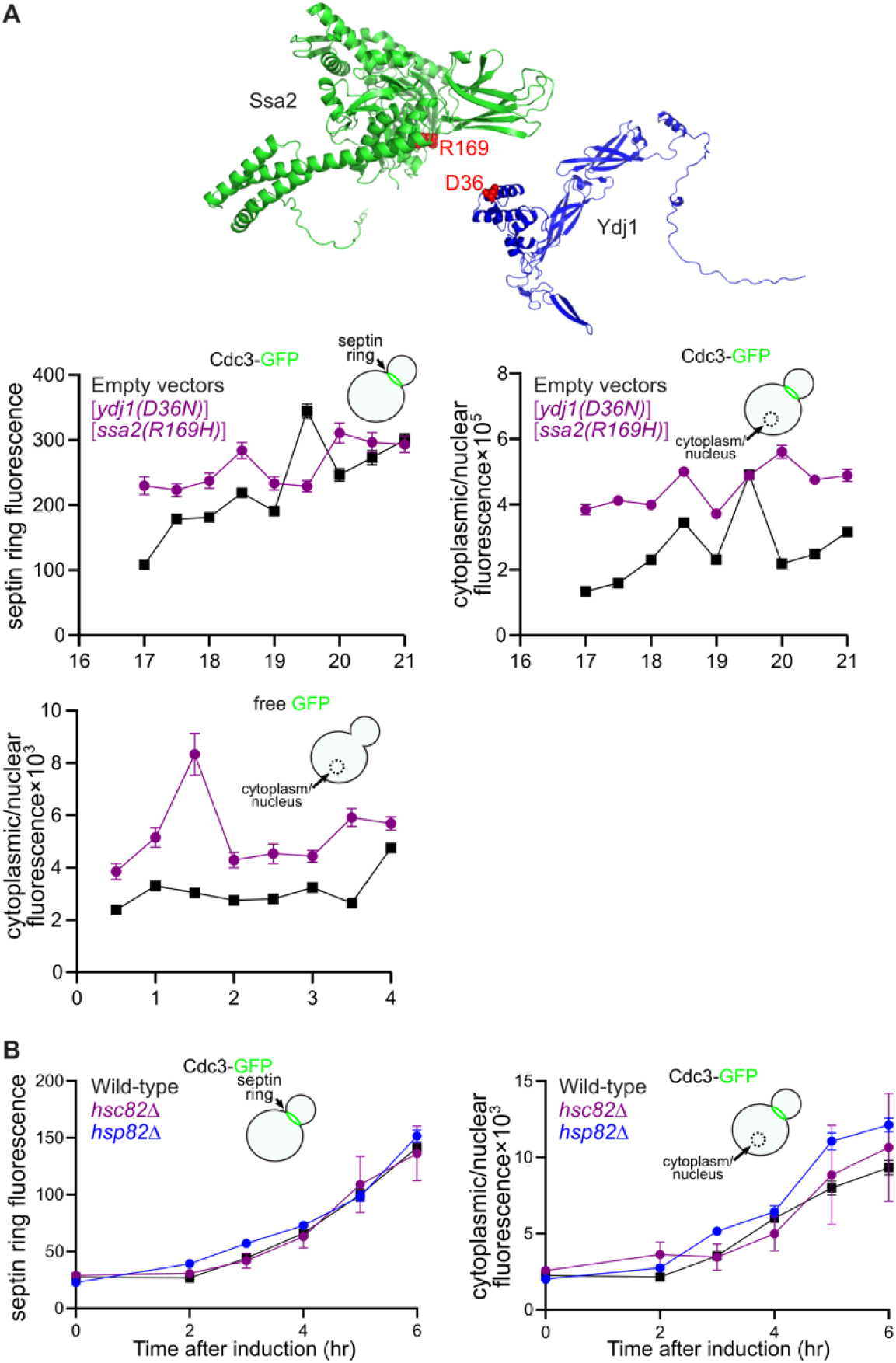
Increased expression from the *GAL1/10* promoter in cells co-expressing an obligatory Hsp40–Hsp70 pair, and lack of effect of cytosolic Hsp80 mutation on the kinetics of septin folding. (A) Above, predicted structures of the cytosolic Hsp70 Ssa2 and the cytosolic Hsp40 Ydj1, showing in red the locations of residues mutated in Ssa2(R169H) and Ydj1(D36N), with the structures juxtaposed approximately how the two proteins are expected to interact. Each mutation inhibits binding to a wild-type partner and restricts binding to the mutant. Plots below show the levels of Cdc3-GFP in yeast septin rings or the cytoplasm/nucleus, or free GFP in the cytoplasm/nucleus, after induction of expression. Yeast cells (strain BY4741) carrying plasmids pMVB1 (“Cdc3-GFP”) or pTS395 (“free GFP”) and pRS314 and pRS315 (“Empty vectors”) or pMR287-D36N and YCpL-SSA2(R169H) (“[*ydj1(D36N)*] [*ssa2(R169H)*]” were grown to mid-log phase in synthetic medium with 2% sodium lactate prior to induction with galactose at final concentration of 0.05% and cells (50 per genotype) from aliquots were imaged at the indicated timepoints. Points show means and error bars are SEM. (B) As in the plots in (A) but cells were carrying only pMVB1; mutant strains were H00418 and H00420.

**Supplemental Figure 4.**
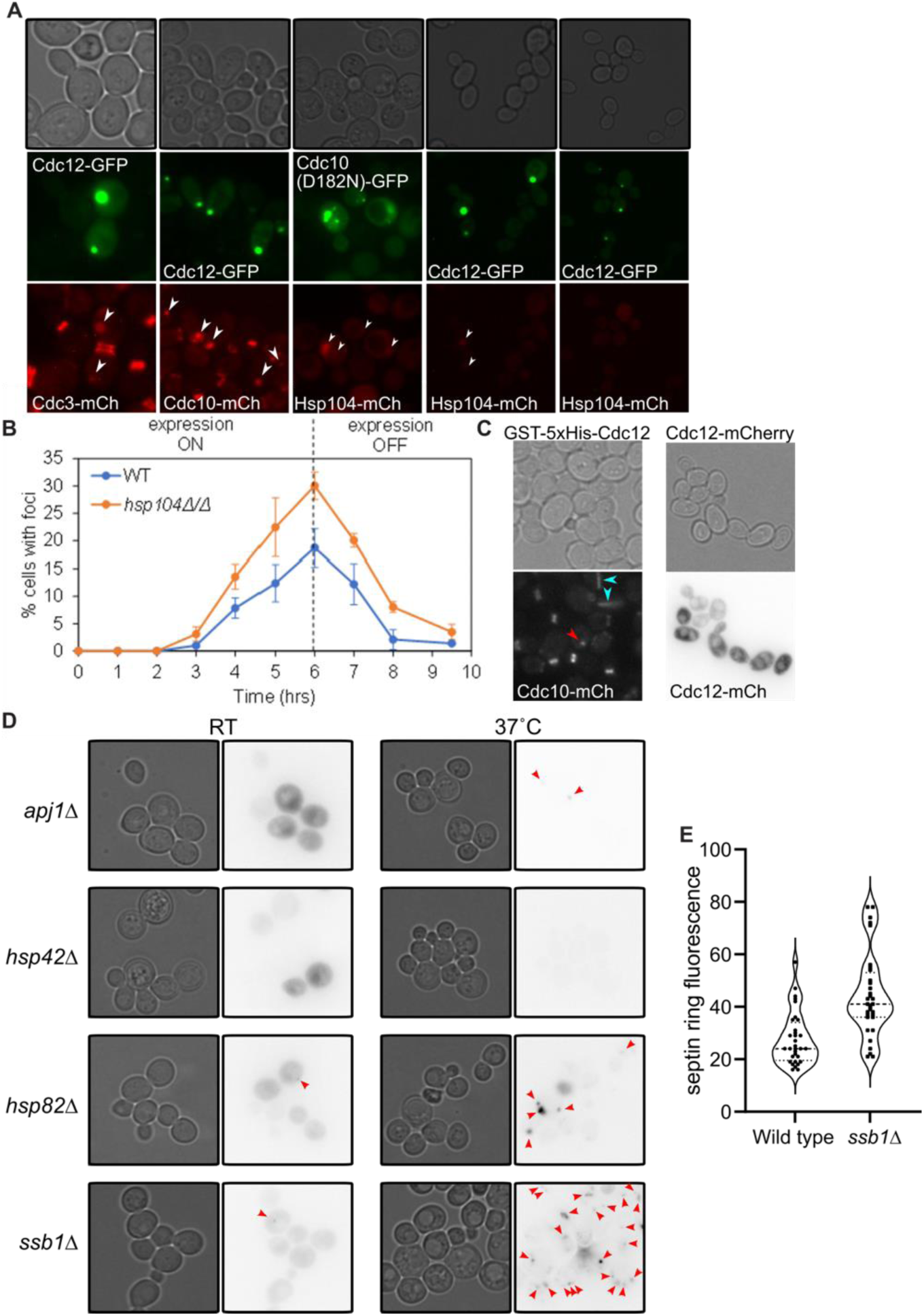
Effects of chaperone mutations and septin overexpression on septin localization. (A) Septin foci and co-localization with other septins and Hsp104. Cells expressing the indicated fluorescently tagged proteins were visualized after overexpression of the GFP-tagged protein from a plasmid (pMVB2 for Cdc12-GFP, pPmet-cdc10-1-GFP for Cdc10(D182N)-GFP). All proteins tagged with mCherry (“mCh”) were encoded at the corresponding chromosomal locus. Strains were BY4743 with integrated *CDC3-mCherry* (“Cdc3-mCh”), JTY3992 (“Cdc10-mCh”), and H06799 (“Hsp104-mCh”). Arrowheads point to foci. (B) Cultures of wild-type (BY4743, “WT”) or *hsp104Δ* diploid cells (strain 31514) carrying plasmid pMVB2 were grown in synthetic medium containing 2% raffinose. At time 0, galactose was added to 0.1% final to induce expression of Cdc12-GFP. After 6 hours, cells were pelleted and resuspended in medium containing glucose and monitored for 3.5 additional hr. At the indicated timepoints, aliquots were removed and imaged and the fraction of cells with foci was calculated from at least 39 cells per genotype per timepoint. Points are means, error bars are standard deviations of the means. (C) As indicated, GST-5xHis-Cdc12 was overexpressed from plasmid YEpPgal–GST-His6-CDC12 in cells of the Cdc10-mCherry-expressing strain JTY3992, or Cdc12-mCherry was overexpressed from plasmid pGF-IVL-470 in strain BY4741. Red arrowhead indicates a focus and cyan arrowheads indicate “rods”. The Cdc12-mCherry image was inverted for clarity. (D) Haploid cells of the indicated genotypes carrying the Cdc10(D182N)-GFP-encoding plasmid pPmet-cdc10-1-GFP were cultured at the indicated temperature (“RT”, room temperature, ∼22°C) in synthetic medium lacking methionine prior to imaging. Fluorescence images were inverted for clarity. Arrowheads point to foci. Strains were H00480, H00500, H00481, and H00483. (E) Haploid cells of the indicated genotype (“WT”, BY4741; “*ssb1Δ*”, Y0059) carrying plasmid pGF-IVL-470 were grown in synthetic medium containing 2% glucose, which represses the *GAL1/10* promoter and should prevent Cdc12-mCherry expression. Signals at septin rings were quantified for 32 cells of each genotype. Within the violin plot, dashed lines indicate the median and dotted lines are quartiles.

**Supplemental Figure 5.**
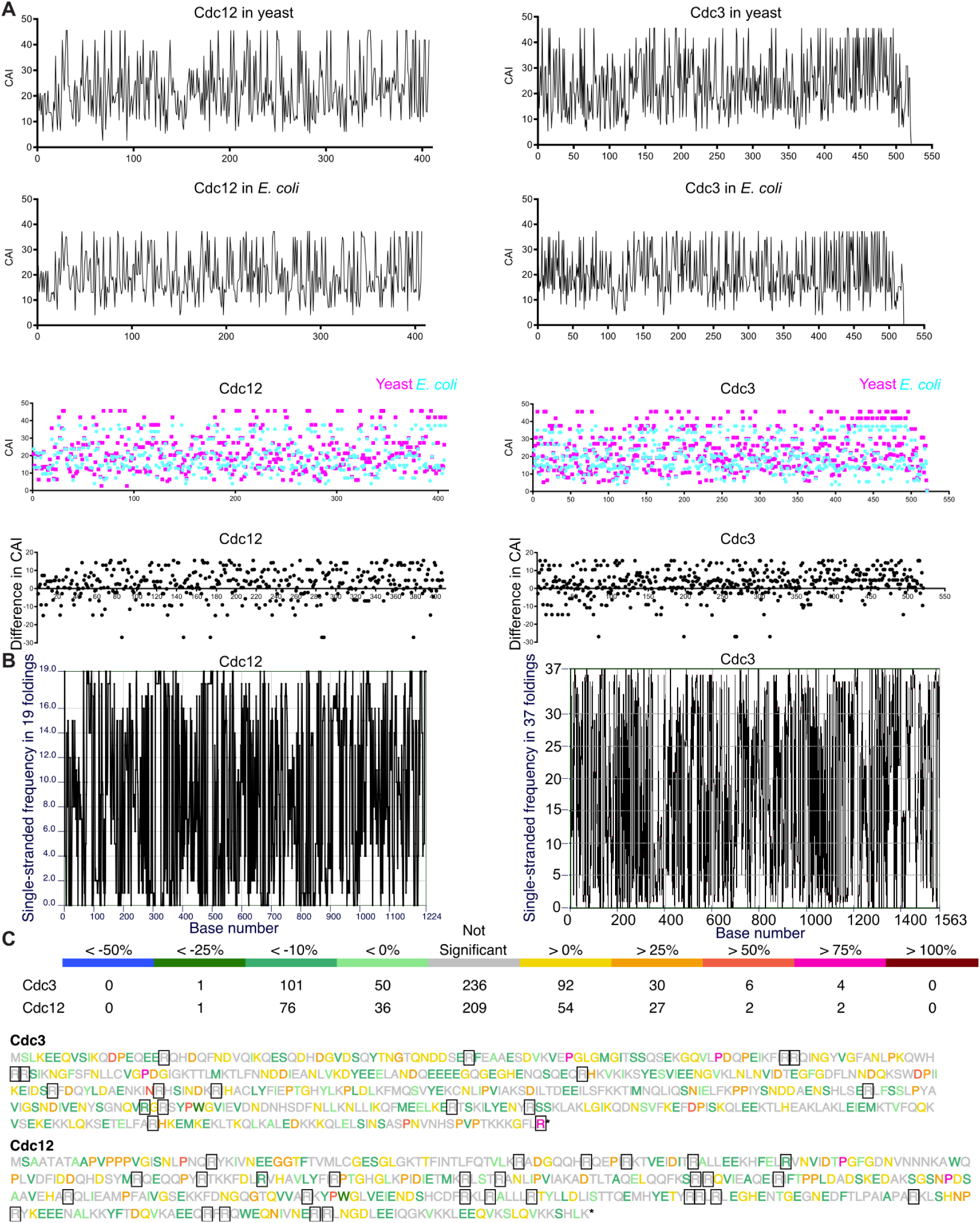
Properties of Cdc3 and Cdc12 that may influence translation elongation. (A) Codon adaptation index (CAI) was calculated for each codon in the Cdc3 and Cdc12 ORFs and plotted against codon position. Top row of plots shows data for *S. cerevisiae* codon usage. Next row shows *E. coli* codon usage, the third row has the same data as the top two but shown as colored points, to highlight regions of mismatch between yeast and *E. coli* that might slow translation specifically in *E. coli*. In the plots in the bottom row, that difference was calculated by subtracting the CAIs from the two species. (B) mRNA secondary structures were predicted using the mFold server and the plots show for each nucleotide in the ORF the number of times it was predicted to be single-stranded. (C) Using bins of semi-quantitative estimates of effects on translation speed (larger values indicated a greater effect on translation) from a published study (Ahmed et al., 2020), each pair of amino acids was analyzed for Cdc3 and Cdc12 and the second amino acid in the pair was assigned a color based on the bin. The total number of pairs in each bin is shown above. Arg residues are boxed.

**Supplemental Figure 6.**
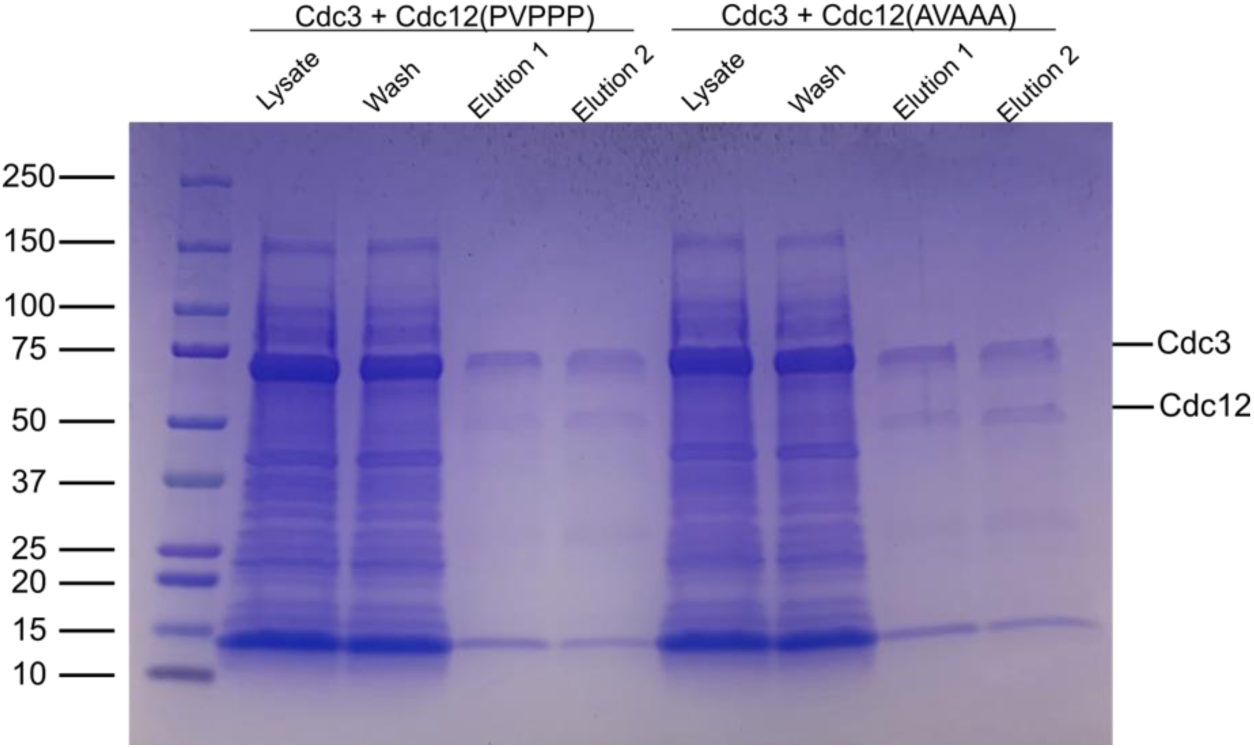
Mutating the Pro-rich cluster at the Cdc12 N terminus does not disrupt folding or expression levels in wild-type *E. coli*. *E. coli* cells (BL21(DE3)) carrying plasmid pMAM54 (“Cdc3 + Cdc12(PVPPP)”) or pMAM88 (“Cdc3 + Cdc12(AVAAA)”) were induced with IPTG, lysed, and the clarified lysate was passed through a Ni-NTA affinity column to purify 6xHis-Cdc12. Aliquots of the lysate, the wash, and the eluate were separated by SDS-PAGE and stained with Coomassie. Leftmost lane contains molecular weight ladder (Bio-Rad #1610395).

**Supplemental Figure 7.**
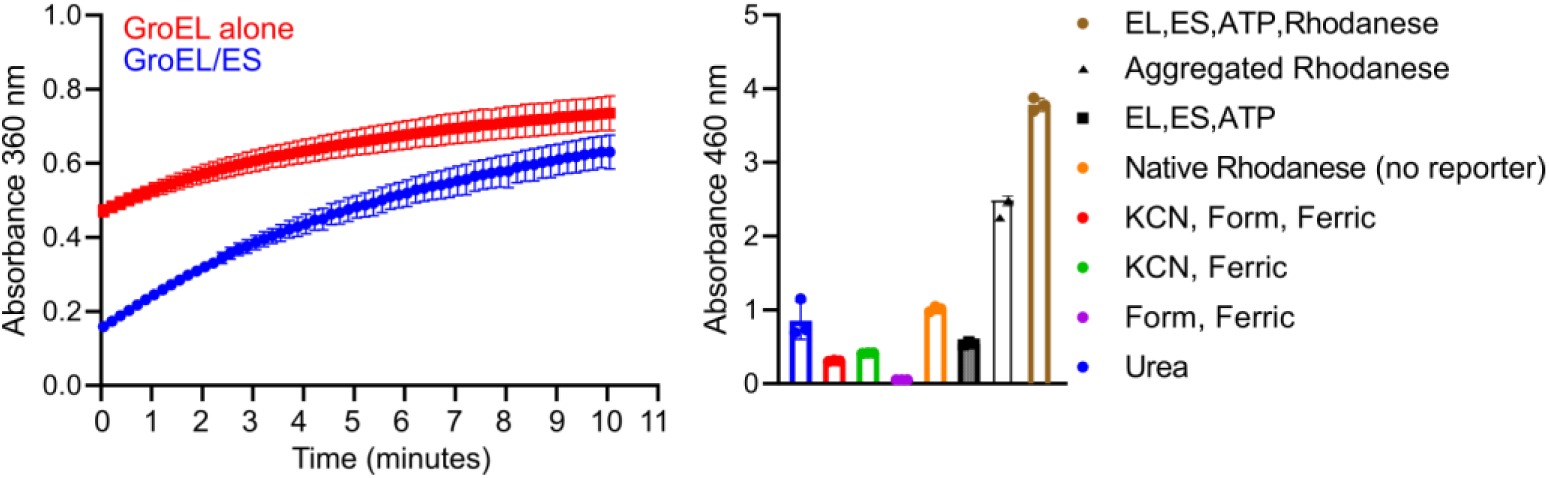
ATPase and refolding activity of purified GroEL/ES. Left, purified GroEL alone or mixed with purified GroES was analyzed with a kit that generates absorbance at 360 nm upon release of inorganic phosphate. The addition of GroES to GroEL is known to inhibit GroEL ATPase activity (Chandrasekhar et al., 1986). Right, rhodanese refolding assay measuring the absorbance at 460 nm of the complex formed between ferric ions and thiocyanate, generated by the activity of rhodanese on thiosulfate and cyanide. Rhodanese was denatured with urea and then diluted into solution containing the “EL”, GroEL; “ES”, GroES; “Form”, formaldehyde, which was added to stop the reactions; “KCN”, potassium cyanide; “Ferric”, ferric nitrate; “no reporter” indicates lack of other reactants. Rhodanese aggregates when it attempts refolding in the absence of GroEL/ES.

**Supplemental Figure 8.**
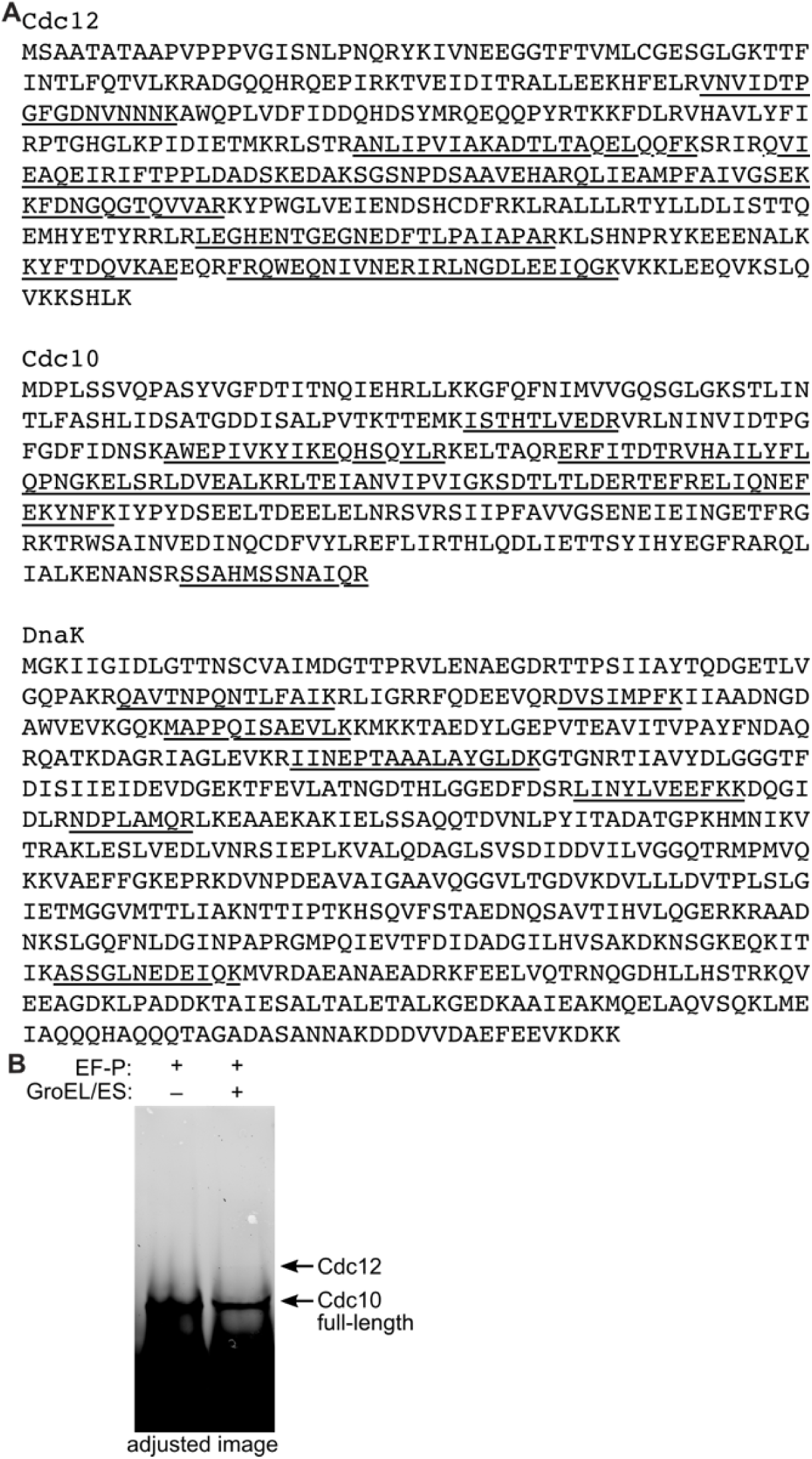
Full-length Cdc12 translated *in vitro* in the presence of GroEL/ES and the lack of effect of added EF-P. (A) Peptides detected by mass spectrometry of an *in vitro* translation reaction performed using template plasmid pMVB128 and in the presence of GroEL/ES are underlined in the protein sequences of Cdc12, Cdc10 and DnaK. (B) As in Figure 5G, but in the presence of added EF-P.

**Supplemental Figure 9.**
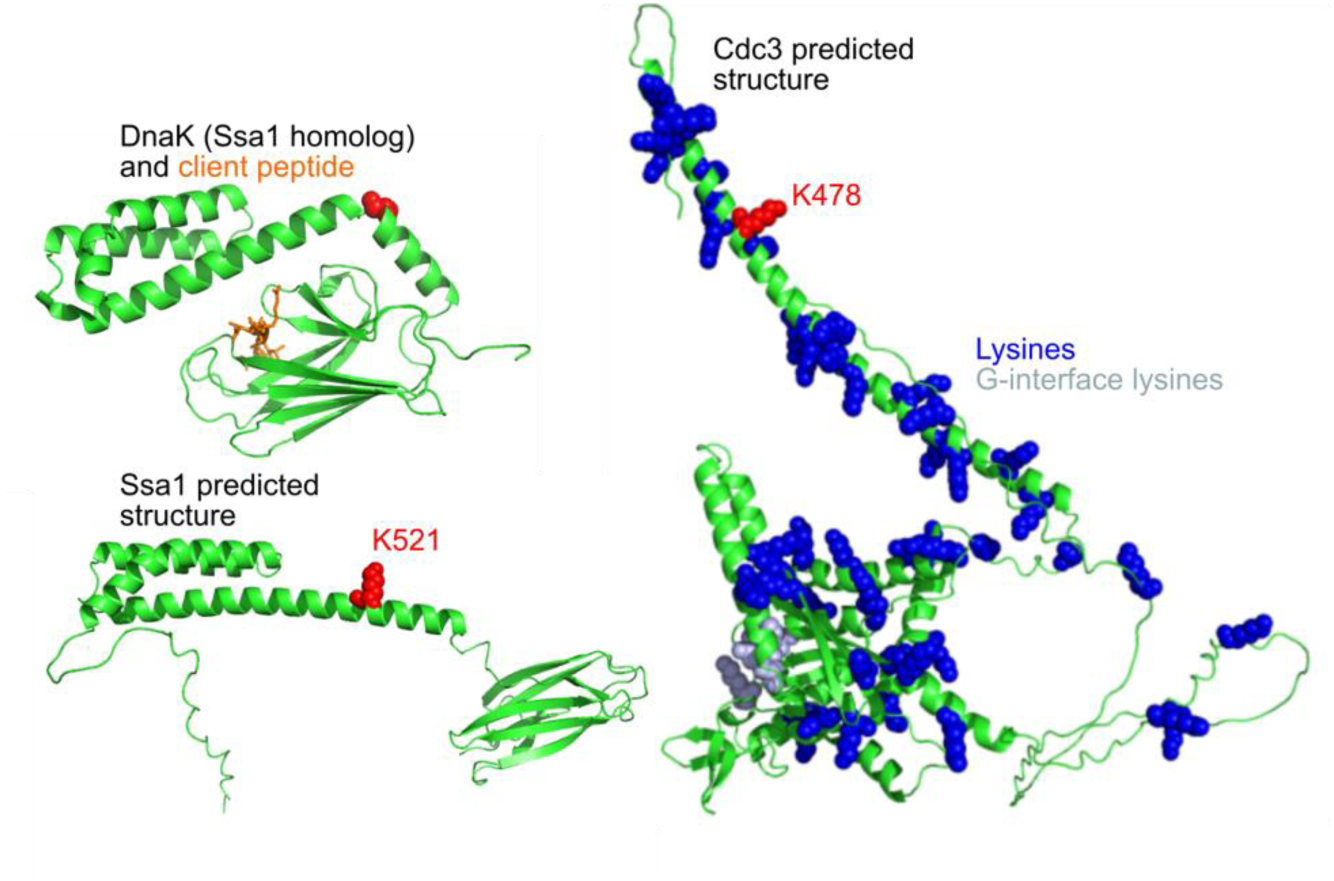
Locations of residues crosslinked between Cdc3 and Ssa1. Structures (solved or predicted) of Cdc3, Ssa1 and the substrate-binding domain of the *E. coli* Hsp70 DnaK bound to a substrate peptide, shown in orange (PDB 1DKZ) (Zhu et al., 1996). Red spheres highlight Lys residues found crosslinked together or, for DnaK, the residue in the position equivalent to Lys521 in Ssa1. In Cdc3, blue spheres highlight all Lys residues, with light blue indicating those in the G interface. Residues 1-394 of Ssa1 were hidden to facilitate comparison with the DnaK structure.

**Supplemental Figure 10.**
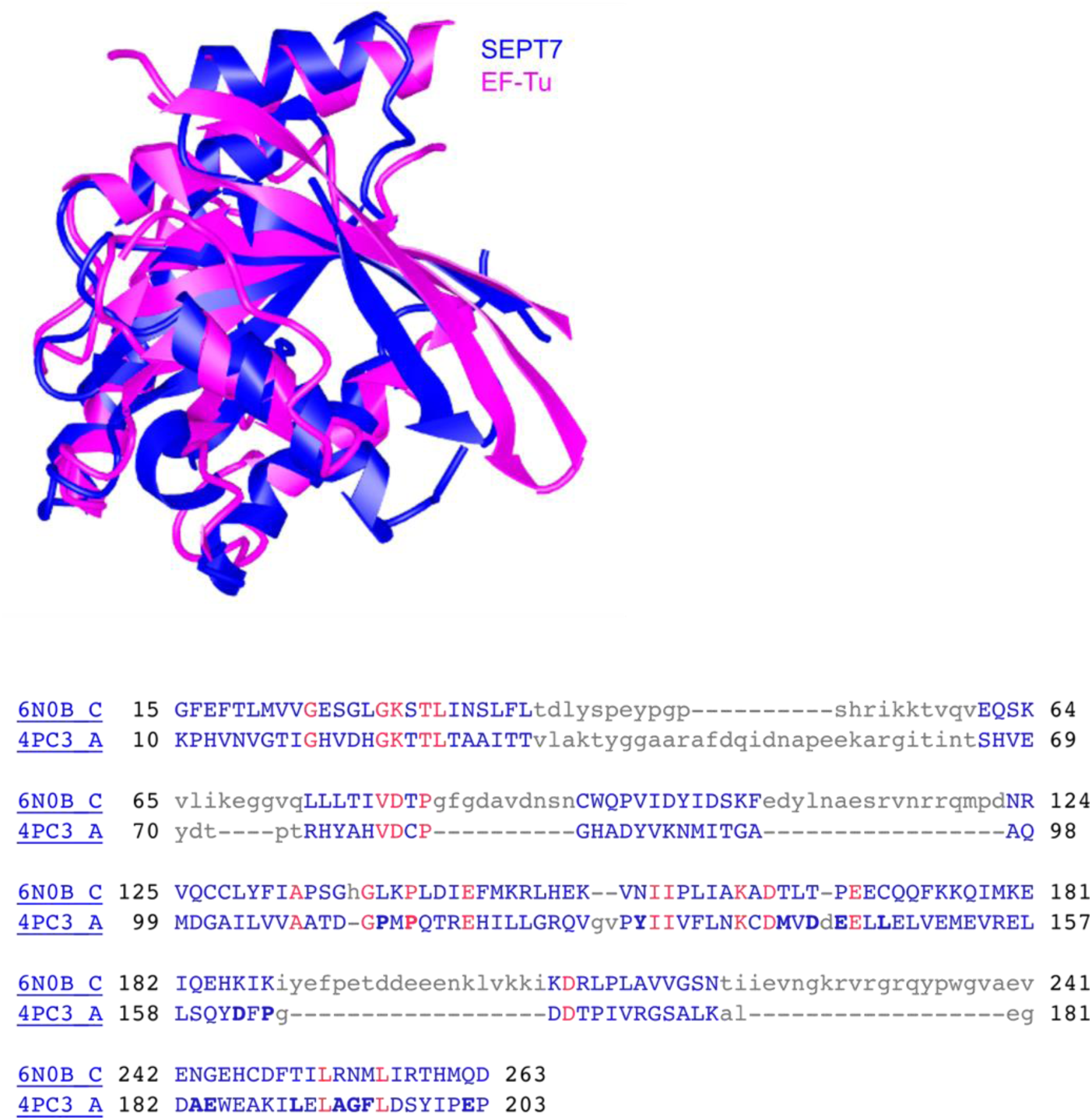
Structural similarities between septins and the GroEL client EF-Tu. Portions of the structure of the GTPase domain of a septin (PDB 6N0B, human septin SEPT7) superimposed on the GTPase domain of EF-Tu (PDB 4PC3). Residues in blue and red in the alignment below are the residues shown in the ribbon structures; red residues are identical between the two proteins.

**Supplemental Figure 11.**
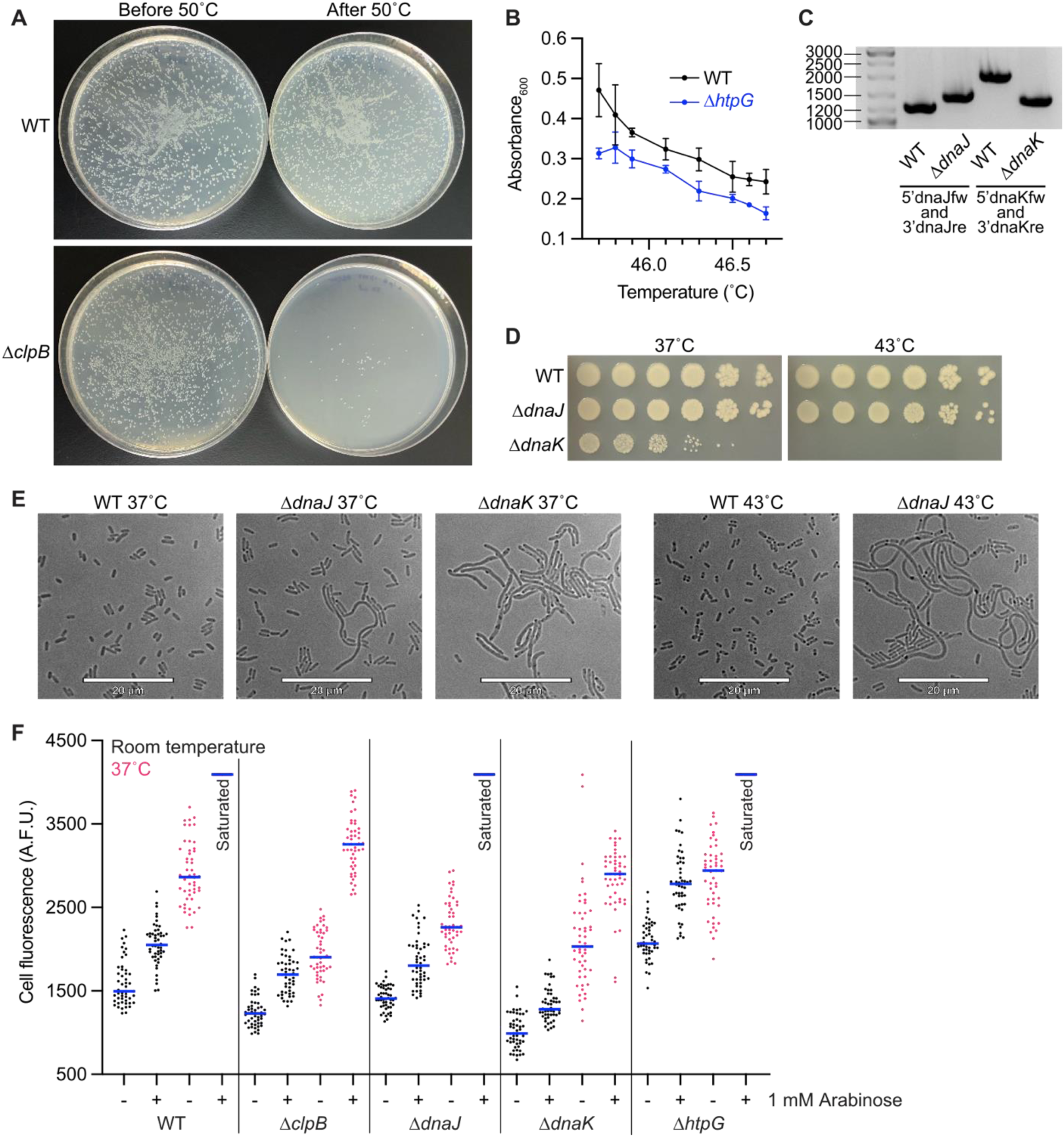
Genotypic and phenotypic characterization of chaperone-deleted and lysogenic strains. (A) Growth of BW25113 (“WT”) and JW2573 (“*ΔclpB*”) cells after 15 min at 42°C and 1 hr at 50°C. 100 µL of a 1000-fold dilution was plated to solid LB medium at 30°C before and after the 50°C incubation for both strains. (B) BW25113 (“WT”) and JW0462 (*ΔhtpG*) cells were grown in 100 µL triplicate cultures in a thermocycler over a 45.7°C to 46.7°C gradient for 6 hrs. Cultures were transferred to a flat-bottom 96-well plate and optical density at 600 nm measured with a Cytation 3 imaging plate reader. Values plotted represent mean and standard deviation as a function of temperature with lines connecting mean values. (C) Genomic DNA was extracted from recombineering candidates B004 (*ΔdnaJ*) and B005 (*ΔdnaK*), in addition to wild-type lysogen B002, and PCR was used to amplify *dnaJ* and *dnaK* loci using 5’dnaJfw and 3’dnaJre or 5’dnaKfw and 3’dnaKre, respectively. (D) Colony growth at 37°C or 43°C by BW25113 (“WT”), B004 (*ΔdnaJ*), and B005 (*ΔdnaK*) cells ten-fold serially diluted in PBS and grown overnight on solid LB medium. (E) BW25113 (“WT”), B004 (*ΔdnaJ*), and B005 (*ΔdnaK*) cells were grown overnight on solid LB medium at the indicated temperatures as in (D) and imaged. (F) Lysogenic strains B002, B003, B004, B005, and B006 carrying T7-driven GFP (pDual-eGFP) were grown to early log phase at 24°C or 37°C in liquid LB medium containing ampicillin. Cells were treated with (1 mM) or without IPTG and grown for another 3 hours at the respective temperature, then imaged. Micrographs were quantified for GFP signal (50 cells for each genotype per temperature and treatment condition). Where >95% of cells for a given treatment produced saturated signal, signal plots were labeled as “saturated.”

